# Mechanisms of Innate Immune Injury in Arrhythmogenic Cardiomyopathy

**DOI:** 10.1101/2023.07.12.548682

**Authors:** Stephen P. Chelko, Vinay Penna, Morgan Engel, Maicon Landim-Vieira, Elisa N. Cannon, Kory Lavine, Jeffrey E. Saffitz

## Abstract

Inhibition of nuclear factor kappa-B (NFκB) signaling prevents disease in *Dsg2^mut/mut^* mice, a model of arrhythmogenic cardiomyopathy (ACM). Moreover, NFκB is activated in ACM patient-derived iPSC-cardiac myocytes under basal conditions *in vitro*. Here, we used genetic approaches and sequencing studies to define the relative pathogenic roles of immune signaling in cardiac myocytes vs. inflammatory cells in *Dsg2^mut/mut^*mice. We found that NFκB signaling in cardiac myocytes drives myocardial injury, contractile dysfunction, and arrhythmias in *Dsg2^mut/mut^* mice. It does this by mobilizing cells expressing C-C motif chemokine receptor-2 (CCR2+ cells) to the heart, where they mediate myocardial injury and arrhythmias. Contractile dysfunction in *Dsg2*^mut/mut^ mice is caused both by loss of heart muscle and negative inotropic effects of inflammation in viable muscle. Single nucleus RNA sequencing and cellular indexing of transcriptomes and epitomes (CITE-seq) studies revealed marked pro-inflammatory changes in gene expression and the cellular landscape in hearts of *Dsg2^mut/mut^* mice involving cardiac myocytes, fibroblasts and CCR2+ cells. Changes in gene expression in cardiac myocytes and fibroblasts in *Dsg2^mut/mut^*mice were modulated by actions of CCR2+ cells. These results highlight complex mechanisms of immune injury and regulatory crosstalk between cardiac myocytes, inflammatory cells, and fibroblasts in the pathogenesis of ACM.

**BRIEF SUMMARY:** We have uncovered a therapeutically targetable innate immune mechanism regulating myocardial injury and cardiac function in a clinically relevant mouse model of Arrhythmogenic Cardiomyopathy (ACM).

## INTRODUCTION

Arrhythmogenic cardiomyopathy (ACM) is a familial heart muscle disease characterized by the early onset of arrhythmias and increased risk of sudden death followed by progressive myocardial injury ultimately leading to heart failure (1–3). Growing evidence implicates inflammation in the pathogenesis of ACM, but mechanisms of immune-mediated injury are not well understood. Inflammation in ACM has usually been considered in the context of inflammatory cell infiltrates in the heart, which occur frequently and can be extensive (3–5). However, although infiltrating inflammatory cells likely contribute to the pathogenesis of ACM (3, 5), no previous studies have validated this conclusion nor have mechanisms of immune cell-mediated myocardial injury been elucidated.

We have reported that signaling mediated via NFκB, a master regulator of the innate immune response (6), is activated in a mouse model of ACM involving homozygous knock-in of a variant in the gene encoding the desmosomal protein, desmoglein-2 (*Dsg2^mut/mut^* mice) (6–8). These mice exhibit clinically relevant features seen in ACM patients, including myocardial injury (cardiac myocyte degeneration and replacement by fibrosis), contractile dysfunction, action potential remodeling and ECG changes, ventricular arrhythmias and inflammation (6–8). Bay 11-7082, a small molecule that inhibits NFκB signaling by blocking degradation of IκBα (9), arrests disease expression and prevents myocardial injury, contractile dysfunction and ECG abnormalities in *Dsg2^mut/mut^* mice (6). NFκB signaling is also activated under basal conditions *in vitro* in iPSC-cardiac myocytes derived from ACM patients with pathogenic variants in *PKP*2(6) or *DSG2* (6, 10). These cells produce large amounts of pro-inflammatory mediators under the control of NFκB without provocation and in the absence of inflammatory cells (6, 10).

The fact that ACM patient iPSC-cardiac myocytes mount an innate immune response and inhibition of NFκB prevents development of disease in *Dsg2^mut/mut^*mice raises a question about the relative pathogenic roles of immune signaling in cardiac myocytes vs. infiltrating inflammatory cells. Here, we answered this question using a genetic approach. To define the role of immune signaling in cardiac myocytes in ACM, we crossed *Dsg2^mut/mut^* mice with a mouse line with cardiac myocyte-specific expression of a dominant-negative form of IκBα (DN-IκBα) (11), which prevents nuclear translocation of NFκB and, thereby, activation of NFκB-mediated changes in gene expression. The resulting double-mutant mice (*Dsg2^mut/mut^* X DN-IκBα) express the *Dsg2* pathogenic variant and can activate NFκB signaling in all cell types except cardiac myocytes. Then, to define the pathogenic role of inflammatory cells in ACM, we focused on cells expressing C-C motif chemokine receptor-2 (CCR2), a G-protein coupled receptor for a monocyte chemo-attractant family that includes monocyte chemoattractant protein-1 (MCP-1, aka CCL2). Cells expressing CCR2 (CCR2+ cells) have been implicated in adverse cardiac remodeling and fibrosis (12, 13). Moreover, MCP-1 expression is increased in hearts of *Dsg2^mut/mut^* mice and in ACM patient iPSC-cardiac myocytes (6). Thus, to define the pathogenic role of CCR2+ cells in ACM, we crossed *Dsg2^mut/mut^* mice with a mouse line with germline deletion of *Ccr2* (*Ccr2^-/-^* mice) to produce double-mutant *Dsg2^mut/mut^* X *Ccr2^-/-^* mice. *Ccr2^-/-^* mice are fertile and have no apparent phenotype under basal conditions, but monocyte/macrophage mobilization in response to immune-activating agents is impaired and macrophages fail to respond to CCL2 (14).

By comparing phenotypes in *Dsg2*^mut/mut^ mice with double-mutant mouse lines (*Dsg2^mut/mut^* X DN-IκB and *Dsg2^mut/mut^* X *Ccr2^-/-^*) and analyzing sequencing data, we observed that activation of NFκB signaling in cardiac myocytes is a major driver of disease in ACM. Myocardial injury and arrhythmias are mediated by CCR2+ inflammatory cells mobilized to the heart via signals from cardiac myocytes. Additionally, CCR2+ cells regulate gene expression in cardiac myocytes and fibroblasts in *Dsg2^mut/mut^* mice. These results highlight complex crosstalk between cardiac myocytes, CCR2+ cells and fibroblasts in immune injury in ACM.

## RESULTS

### NFκB signaling in cardiac myocytes leads to myocardial injury, contractile dysfunction and arrhythmias in *Dsg2^mut/mut^* mice and mobilizes CCR2+ cells to the heart

To define the role of innate immune signaling via NFκB in cardiac myocytes in the pathogenesis of ACM, we compared phenotypes of 16-week-old WT, *Dsg2^mut/mut^* and *Dsg2^mut/mut^* X DN-*IĸBα*^+/-^ mice. As reported previously (6, 7), 16-week old *Dsg2*^mut/mut^ mice show considerable myocardial injury indicated by the presence of extensive fibrosis (which had replaced areas of degenerated cardiac myocytes), marked contractile dysfunction with significantly reduced left ventricular ejection fraction, numerous PVCs and alterations in signal-averaged ECGs (SAECGs) when compared to age-matched WT mice (**Figure 1**, **Table 1**). All of these key disease features were markedly attenuated in 16-week-old *Dsg2^mut/mut^*X DN-*IĸBα*^+/-^ mice (**Figure 1**, **Table 1**). These results indicate that activation of NFκB signaling in cardiac myocytes leads to myocardial injury, contractile dysfunction and arrhythmias in *Dsg2*^mut/mut^ mice.

**Figure 1.**
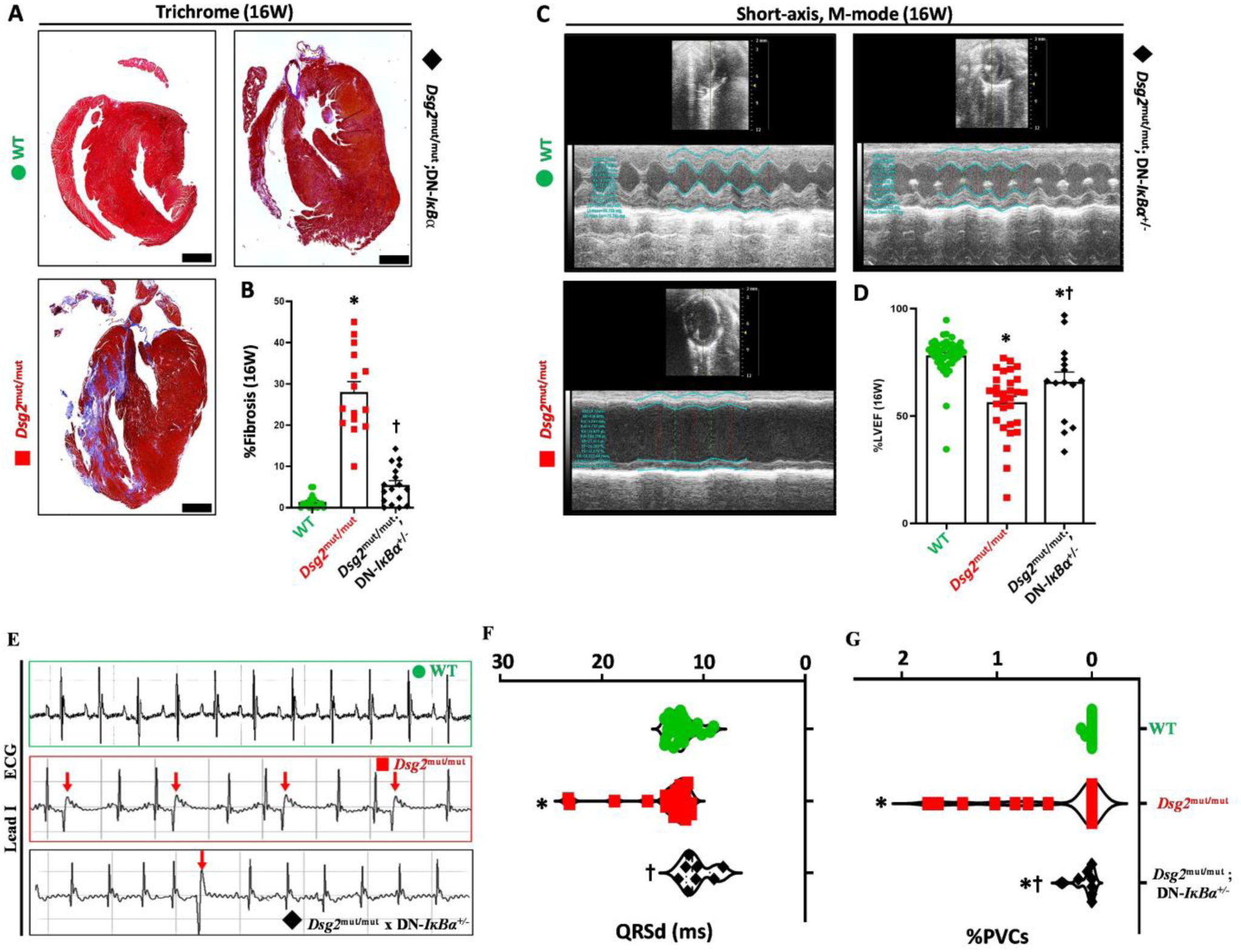
Blocking activation of NFĸB signaling in cardiac myocytes mitigates myocardial injury and arrhythmias and preserves cardiac function in *Dsg2^mut/mut^* mice. (**A**) Representative images of trichrome-stained hearts from mice at 16 weeks of age (16W) selected from ≥10 mice/cohort/time-point. (**B**) Percent (%) fibrosis at 16W; scale bar, 1mm. (**C**) Representative echocardiographs from WT, *Dsg2^mut/mut^*, and *Dsg2^mut/mut^* X DN-*IĸBα^+/-^* mice at 16W, selected from ≥10 mice/cohort/time-point. (**D**) Percent left ventricular ejection fraction (%LVEF). Note preserved cardiac function in *Dsg2^mut/mut^* X DN-*IĸBα^+/-^* mice. * P<0.001 any cohort vs WT; ^†^ P<0.001 any cohort vs *Dsg2^mut/mut^*using one-way ANOVA with Tukey’s posthoc analysis. (**E**) Representative ECGs from WT, *Dsg2*^mut/mut^ and *Dsg2*^mut/mut^ X DN-*IĸBα^+/-^* mice at 16W. Premature ventricular contractions (PVCs) are noted by red arrows. (**F, G**) QRS duration (QRSd) and percent PVCs (%PVCs), respectively, obtained from signal averaged ECGs. * P<0.05 any cohort vs WT; ^†^ P<0.05 any cohort vs *Dsg2*^mut/mut^; using one-way ANOVA with Tukey’s posthoc analysis.

**Table 1.** Morphometric and electrocardiographic data at 16 weeks of age.

| Parameter | WT | *Dsg2*<sup>mut/mut</sup> | *Dsg2*<sup>mut/mut</sup><br>x *Ccr2*<sup>-/-</sup> | *Dsg2*<sup>mut/mut</sup><br>x DN-*IkBa*<sup>+/-</sup> |
| --- | --- | --- | --- | --- |
| **Morphometric** |  |  |  |  |
| LVM (mg) | 82.9 ± 3.3 | 96.4 ± 3.5* | 108.4 ± 10.4*<sup>‡</sup> | 79.9 ± 5.0<sup>†</sup> |
| HW/BW (mg/g) | 4.65 ± 0.1 | 5.31 ± 0.2* | 4.99 ± 0.1* | 4.88 ± 0.1<sup>†</sup> |
| **Electrocardiography** |  |  |  |  |
| Heart Rate (bpm) | 486 ± 9 | 495 ± 7 | 440 ± 29*<sup>†</sup> | 471 ± 22 |
| RR-I (ms) | 125 ± 2.2 | 122 ± 1.7 | 140 ± 8.9*<sup>†</sup> | 131 ± 6.9<sup>†</sup> |
| PR-I (ms) | 39.6 ± 0.8 | 40.5 ± 1.1 | 40.9 ± 2.5 | 44.7 ± 3.4* |
| Pd (ms) | 10.5 ± 0.3 | 10.5 ± 0.5 | 12.1 ± 0.7* | 12.1 ± 0.8* |
| QRSd (ms) | 12.2 ± 0.3 | 13.8 ± 0.7* | 14.1 ± 1.0*<sup>‡</sup> | 11.7 ± 0.5<sup>†</sup> |
| P-Amp (mV) | 0.07 ± 0.004 | 0.05 ± 0.01* | 0.06 ± 0.01 | 0.05 ± 0.01* |
| R-Amp (mV) | 0.73 ± 0.1 | 0.64 ± 0.1 | 0.57 ± 0.1 | 0.64 ± 0.1 |
| Q-Amp (mV) | -0.05 ± 0.01 | -0.09 ± 0.01* | -0.06 ± 0.02 | -0.07 ± 0.02 |
| S-Amp (mV) | -0.23 ± 0.02 | -0.10 ± 0.03* | -0.14 ± 0.05 | -0.04 ± 0.04* |

WT, wildtype; LVM, left ventricular mass; HW, heart weight; BW, body weight; RR-I, R-R interval; PR-I, P-R interval; Pd, P-wave duration; QRSd, QRS-wave duration; P-Amp, P-wave amplitude; R-Amp, R-wave amplitude; Q-Amp, Q-wave amplitude; S-Amp, S-wave amplitude. PVCs, premature ventricular contractions; Data presented as mean±SEM, n≥15mice/cohort/parameter. \*P<0.05 any cohort vs WT; <sup>†</sup>P<0.05 any cohort vs *Dsg2*<sup>mut/mut</sup> mice; <sup>‡</sup>P<0.05 *Dsg2*<sup>mut/mut</sup> x *Ccr2*<sup>-/-</sup> vs *Dsg2*<sup>mut/mut</sup> x DN-*IkBa*<sup>+/-</sup> using One-way ANOVA with Tukey's post-hoc test.

To determine if activation of NFκB signaling in cardiac myocytes also affects myocardial inflammatory cells in ACM, we measured numbers of macrophage subsets in hearts of 16-week-old *Dsg2^mut/mut^* and *Dsg2^mut/mut^* X DN-*IĸBα*^+/-^ mice. We focused on macrophages because they are the most abundant immune cell type in mouse and human hearts (15). As shown in **Figure 2**, the number of CD68+ macrophages was greatly increased in hearts in *Dsg2^mut/mut^* compared to WT mice. Furthermore, the percentage of total myocardial CD68+ cells expressing Lyve1 (lymphatic vessel endothelial hyaluronic acid receptor 1), a marker of cardiac resident macrophages (16, 17), was substantially diminished in *Dsg2^mut/mut^* mice reflecting a shift in macrophage ontogeny favoring recruited CCR2+ monocyte-derived macrophages. While the total number of myocardial CD68+ macrophages was equivalent in *Dsg2^mut/mut^* and *Dsg2^mut/mut^*X DN-*IĸBα*^+/-^ mice, macrophage populations in *Dsg2^mut/mut^* X DN-*IĸBα*^+/-^ hearts were more reminiscent of those seen in hearts of WT animals. Specifically, *Dsg2^mut/mut^* X DN-*IĸBα*^+/-^ mice contained fewer CCR2+ monocyte-derived macrophages and increased numbers of Lyve1+ macrophages compared to *Dsg2^mut/mut^* hearts. Taken together, these results indicate that NFκB signaling in cardiac myocytes leads to accumulation of CCR2+ macrophages and loss of cardiac resident macrophages. Such an alteration in cardiac macrophage compositional shift has been associated with enhanced myocardial inflammation and reduced capacity for tissue repair (16).

**Figure 2.**
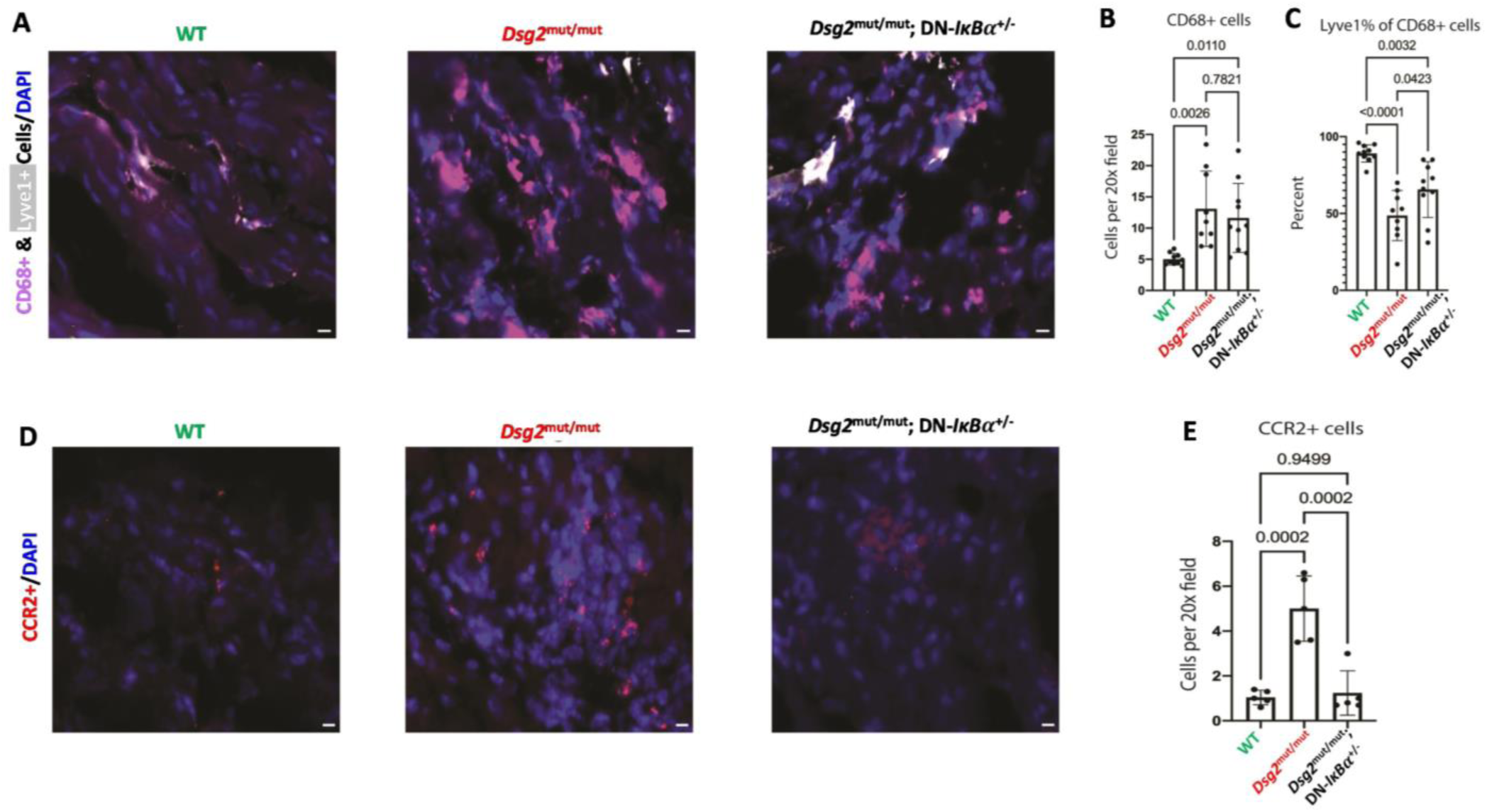
NFкB signaling in cardiac myocytes mobilizes inflammatory cells to the heart in *Dsg2*^mut/mut^ mice. (**A**) Representative immunofluorescence staining of myocardial sections from 16-week-old (16W) mice showing CD68+ (violet) and Lyve1+ (white) cells. (**B, C**) Quantification of CD68+ cells and Lyve1+ cells as a percentage of CD68+ cells in mice. (**D**) Representative RNA *in situ* hybridization images (RNAscope) showing CCR2+ (red) cells in 16-week-old mice. (**E**) Quantification of CCR2+ cells in mice of the indicated genotypes. P-values determined by one-way ANOVA; WT, n=10 samples; *Dsg2^mut/mut^*, n=9 samples; *Dsg2^mut/mut^*; DN-*IĸBα^+/-^*, n=10 samples.

### CCR2+ cells promote myocardial injury and arrhythmias in *Dsg2^mut/mut^*mice

Deletion of *Ccr2* impairs monocyte egress from the bone marrow and spleen, and recruitment of monocytes to peripheral sites (14, 18). Accordingly, to define the role of monocyte recruitment in the pathogenesis of ACM, we compared phenotypes in 16-week-old WT, *Dsg2^mut/mut^* and *Dsg2^mut/mut^* X *Ccr2*^-/-^ mice. While the number of myocardial CD68+ cells was increased in both *Dsg2^mut/mut^* and *Dsg2^mut/mut^* X *Ccr2*^-/-^ mice compared to WT, the percentage of Lyve1+ cardiac resident macrophages was equivalent in *Dsg2^mut/mut^* X *Ccr2*^-/-^ and WT hearts (**Figure 3**). As also shown in **Figure 3**, the amount of myocardial fibrosis in *Dsg2^mut/mut^* X *Ccr2*^-/-^ mice was markedly less than that seen in *Dsg2^mut/mut^*mice. *Dsg2^mut/mut^* X *Ccr2*^-/-^ mice also showed a significant reduction in the number of PVCs and less severe alterations in SAECGs compared to *Dsg2^mut/mut^* mice. These results indicate that monocyte recruitment and the resulting accumulation of CCR2+ macrophages mediate myocardial injury/fibrosis and promotes arrhythmias in *Dsg2^mut/mut^* mice. They also suggest that the principal mechanism by which NFκB signaling in cardiac myocytes drives myocardial injury and arrhythmias is by mobilizing CCR2+ cells to the heart where they act as injurious agents. Even so, despite the significant reduction in myocardial fibrosis in *Dsg2^mut/mut^* X *Ccr2*^-/-^ mice, there was no apparent improvement in left ventricular ejection fraction. The degree of contractile dysfunction in *Dsg2^mut/mut^*X *Ccr2*^-/-^ mice was equivalent to that seen in age-matched *Dsg2^mut/mut^* mice (**Figure 3**).

**Figure 3.**
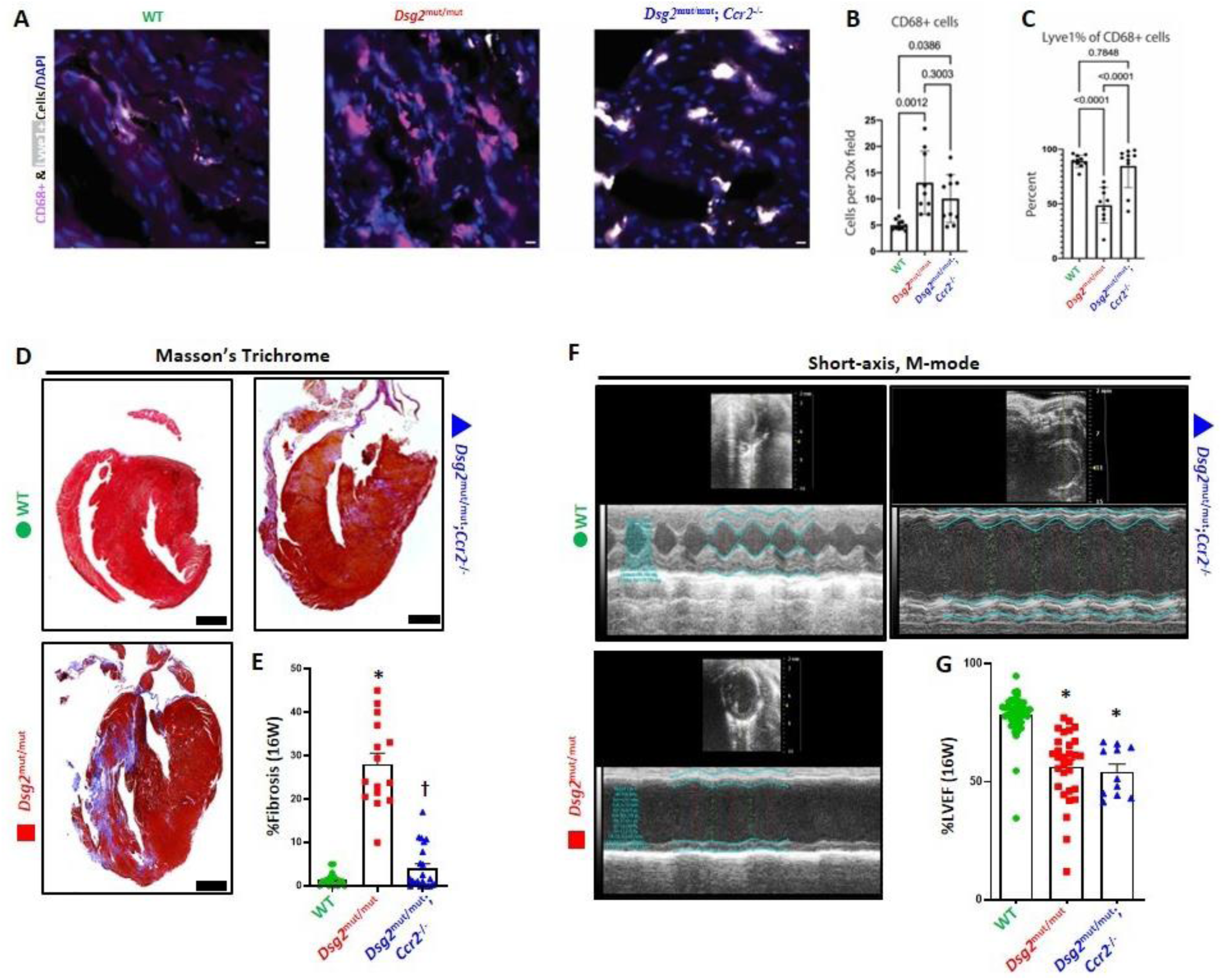
CCR2+ cells contribute to myocardial injury and dysfunction. (**A**) Representative myocardial images depicting CD68+ (violet) and Lyve1+ (white) cells via immunofluorescence staining from 16W old mice. (**B, C**) Quantification of CD68+ cells and Lyve1+ cells as a percentage of CD68+ cells in mice. (**D**) Representative trichrome-stained hearts from 16-week-old mice. Images were selected from ≥10mice/cohort/time-point. Black scale bar, 1mm. (**E**) Percent (%) fibrosis at 16 weeks. (**F**) Representative echocardiograms from WT, *Dsg2^mut/mut^*, and *Dsg2^mut/mut^* X *Ccr2*^-/-^ mice. (**G**) Percent left ventricular ejection fraction (%LVEF). Note, reduced fibrosis in *Dsg2^mut/mut^* X *Ccr2*^-/-^ mice but without preserved cardiac function. *P<0.001 any cohort vs WT; ^†^ P<0.001 any cohort vs *Dsg2^mut/mut^* using One-way ANOVA with Tukey’s posthoc analysis.

### NFκB signaling in cardiac myocytes and CCR2+ cells both contribute to production of inflammatory mediators in *Dsg2*^mut/mut^ mice

Levels of pro-inflammatory cytokines and fibrokines are increased in hearts of 16-week-old *Dsg2^mut/mut^* mice (6). These mediators are also produced under basal conditions *in vitro* by cardiac myocytes derived from iPSCs from ACM patients with variants in *PKP2* or *DSG2* (6, 10). To gain insights into the sources of inflammatory mediators in ACM, we compared cytokine levels in hearts of 16-week-old WT, *Dsg2^mut/mut^* and double-mutant mice. As shown in **Figure 4**, levels of major cytokines of the innate immune response including IL-1β, IL-6 and MCP-1 (aka, CCL2) were increased in hearts of 16-week *Dsg2^mut/mut^* mice compared to WT mice (complete cytokine expression data are shown in **Supplemental Table 1**). Nearly all of these inflammatory mediators were at least partially normalized in hearts of either *Dsg2^mut/mut^*X DN-*IĸBα*^+/-^ and *Dsg2^mut/mut^* X *Ccr2*^-/-^ mice (**Figure 4** and **Supplemental Table 1**), suggesting that cardiac myocyte NFκB signaling and CCR2+ macrophages both trigger expression of inflammatory mediators in *Dsg2^mut/mut^* mice. One exception was osteopontin (OPN) which was greatly increased in hearts of *Dsg2^mut/mut^* and *Dsg2^mut/mut^* X *Ccr2*^-/-^ mice, but not in *Dsg2^mut/mut^* X DN-*IĸBα*^+/-^ mice (**Figure 4**). Of note, the fibrosis marker periostin (POSTN) (19, 20) was greatly increased in *Dsg2^mut/mut^*hearts, but not in either *Dsg2^mut/mut^* X DN-*IĸBα*^+/-^ or *Dsg2^mut/mut^*X *Ccr2*^-/-^ hearts, both of which showed significantly reduced amounts of ventricular fibrosis. To gain further insights into the contribution of monocyte recruitment and monocyte-derived CCR2+ macrophages in the pathogenesis of ACM and identify cell sources of cytokine production, we utilized established single cell transcriptomic pipelines.

**Figure 4.**
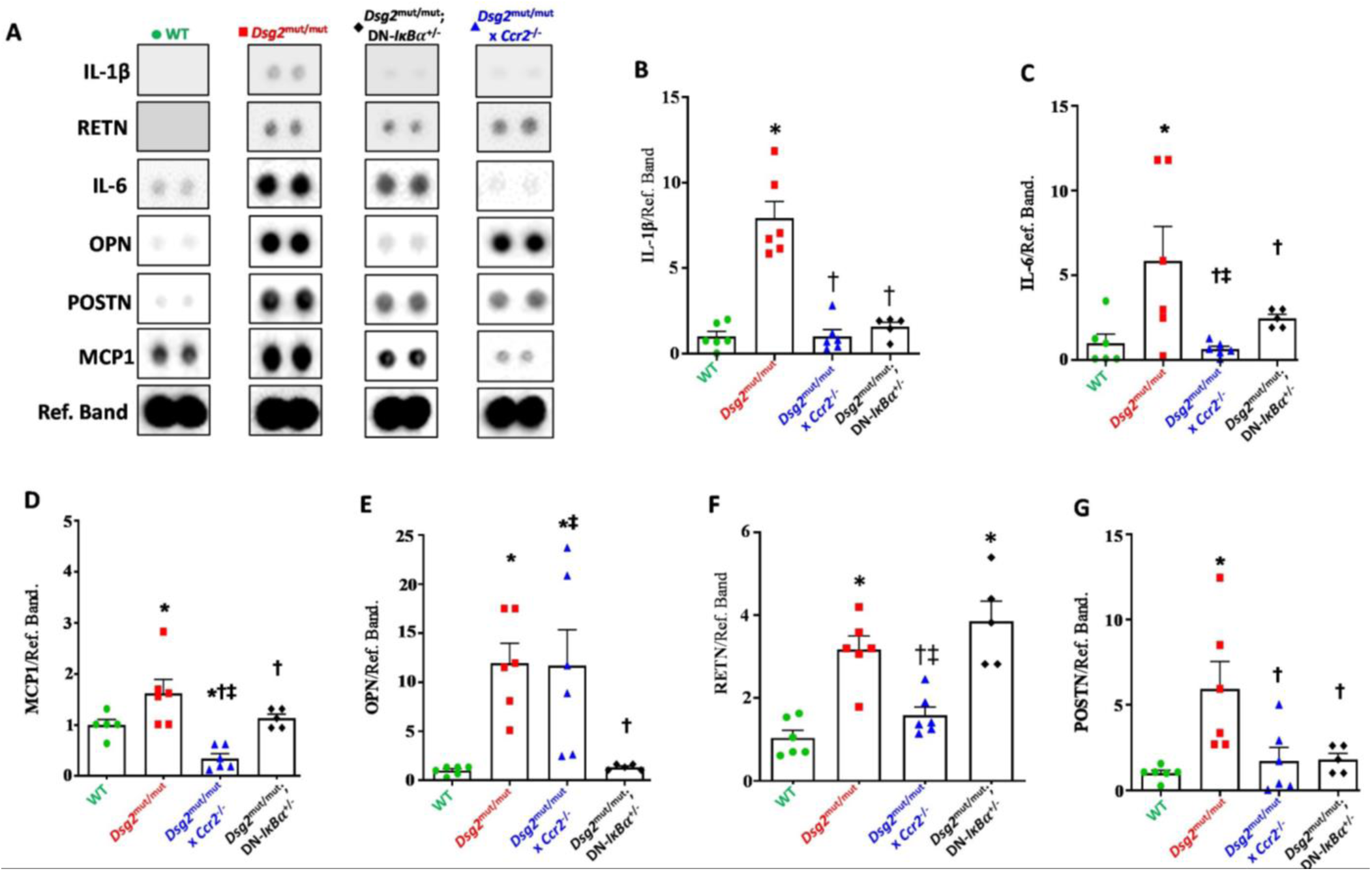
NFκB signaling in cardiac myocytes and actions of CCR2+ cells increase myocardial cytokines levels in *Dsg2^mut/mut^* mice. (**A**) Representative immunoblots from myocardial cytokine arrays in WT, *Dsg2^mut/mut^*, *Dsg2^mut/mut^ X Ccr2^-/-^* and *Dsg2^mut/mut^* X DN-*IĸBα^+/-^* mice. Ref. Band, Reference Band. (**B-G**) Bar graphs comparing levels of selected cytokines with levels in WT hearts normalized to 1. Data presented as mean±SEM; *P<0.05 any cohort vs WT; ^†^ P<0.05 any cohort vs *Dsg2^mut/mut^*; ^‡^P<0.05 any cohort vs *Dsg2^mut/mut^* X DN-*IĸBα^+/-^* using one-way ANOVA with Tukey’s posthoc analysis.

### CITE-seq reveals expansion of CCR2+ inflammatory macrophages in hearts of *Dsg2^mut/mut^* mice

To characterize the transcriptional and cell surface proteomic landscape in ACM, we performed CITE-seq on pooled hearts from 16-week-old WT, *Dsg2^mut/mut^* and *Dsg2^mut/mut^* X *Ccr2*^-/-^ mice (**Figure 5A**). After pre-processing and application of quality control filters (**Supplemental Figure 1A-C**) (15, 21), we identified 7 distinct stromal and immune cell types (**Figure 5B, Supplemental Figure 1D, E**) including fibroblasts, endothelial cells, pericytes/smooth muscle cells, monocytes/macrophages, neutrophils and T cells. Differential gene expression analysis revealed cell-type specific transcriptional differences across all major cell types in WT vs. *Dsg2^mut/mut^* and *Dsg2^mut/mut^* vs *Dsg2^mut/mut^* X *Ccr2*^-/-^ conditions, which were especially prominent in monocytes/macrophages and fibroblasts (**Supplemental Figure 1F, G**). Given the prominent inflammation and fibrosis observed in *Dsg2*^mut/mut^ mice, we focused on monocyte/macrophage and fibroblast populations. Within the monocyte/macrophage cluster, we identified several subpopulations including Lyve1+ macrophages, Trem2+ macrophages, type-1 and -2 conventional dendritic cells, CCR2+ monocytes and macrophages, and Ly6c low monocytes (**Figure 5C, Supplemental Figure 2A, B**). Consistent with the immunostaining and *in situ* hybridization studies presented above, cell composition and kernel density analysis revealed increased proportions of CCR2+ monocytes and macrophages and decreased Lyve1+ macrophages in *Dsg2^mut/mut^* hearts compared to WT hearts. These populations were restored to WT levels in *Dsg2^mut/mut^* X *Ccr2*^-/-^ hearts (**Figure 5D, E**). Differential gene expression analysis between WT vs *Dsg2^mut/mut^* monocytes/macrophages revealed profound differences with increased expression of many inflammatory and fibrosis-associated genes (e.g., *Plac8, Ly6c2, Ccl6, Lgals3*)(16) and decreased expression of resident macrophage-associated genes (e.g., *Cd163, Mrc1, Folr2*) (16, 17) in *Dsg2^mut/mut^* hearts. These gene expression changes were normalized in *Dsg2^mut/mut^* X *Ccr2*^-/-^ mice (**Figure 5F, H**). Projection of genes differentially expressed by monocytes/macrophages in *Dsg2*^mut/mut^ hearts within the UMAP space revealed that CCR2+ monocytes and macrophages were the primary source of inflammatory mediators enriched in *Dsg2^mut/mut^*hearts (**Figure 5G**). Pathway enrichment analysis indicated that genes differentially expressed in *Dsg2^mut/mut^* monocytes/macrophages were associated with increased innate immune activation and fibroblast proliferation (**Figure 5I**).

**Figure 5.**
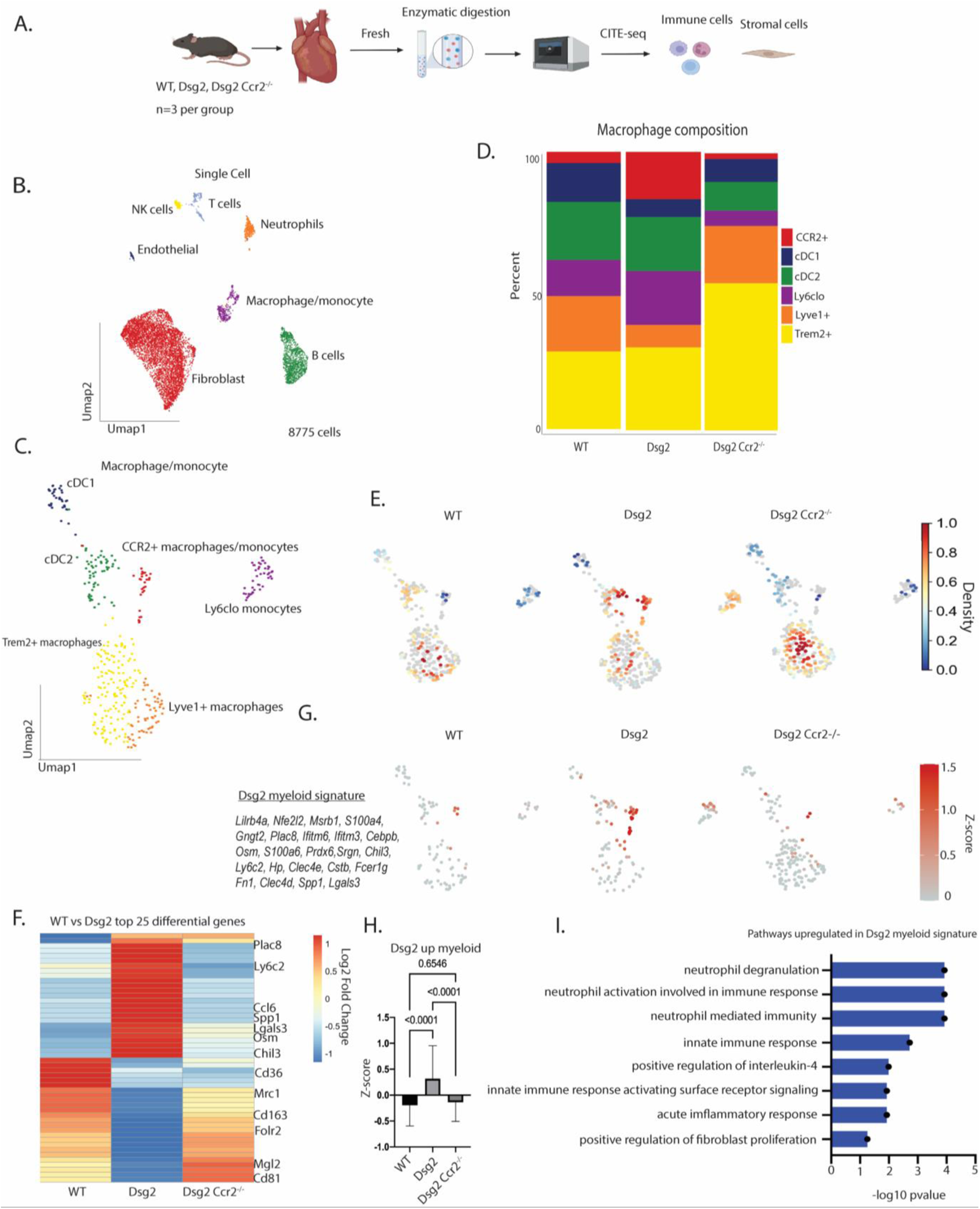
CITE-seq reveals expansion of CCR2+ inflammatory macrophages in hearts of *Dsg2^mut/mut^* mice. **(A)** Schematic depicting design of CITE-seq experiments. Whole hearts were homogenized and enzymatically digested. **(B)** Uniform Manifold Approximation and Projection (UMAP) of 8,775 cells after QC and data filtering using standard Seurat pipeline. **(C)** Uniform Manifold Approximation and Projection (UMAP) re-clustering of macrophage/monocyte population. **(D)** Composition graph showing proportion of different populations within the macrophage/monocyte cluster. **(E)** Gaussian kernel density estimation of cells within the macrophage/monocyte cluster across the indicated genotypes. **(F)** Heatmap of top 25 differentially expressed genes in the macrophage/monocyte cluster between WT and *Dsg2^mut/mut^* mice with side-by-side comparison of the expression of those same genes from *Dsg2^mut/mut^* X *Ccr2^-/-^* mice. Example genes are annotated. (**G**) Z-score feature plot, overlaying an inflammatory gene signature derived from the heatmap in **F** (genes listed to the side) and displayed on the macrophage/monocyte UMAP projection split by genotype. (**H**) Graph of z-score values for inflammatory gene signature derived from heatmap in **F** compared across genotypes (P values determined by one-way ANOVA test). (**I**) Top KEGG pathways for top 25 differentially upregulated genes in *Dsg2*^mut/mut^ mice (derived from **F**).

### *Postn+* fibroblasts are expanded in *Dsg2^mut/mut^* hearts through a CCR2+ monocyte- and macrophage-dependent mechanism

To determine how inhibition of CCR2+ monocyte recruitment impacts cardiac fibroblasts in ACM, we further analyzed the fibroblast cluster and identified a number of transcriptionally distinct fibroblast states (**Figure 6a, Supplemental Figure 3A, C**). Cell composition and kernel density analysis revealed a profound increase in *Postn*+ fibroblasts and a decrease in *Cxcl14+* fibroblasts in *Dsg2^mut/mut^* hearts compared to WT hearts (**Figure 6B, Supplemental Figure 3B**). We next performed differential gene expression analysis in WT and *Dsg2*^mut/mut^ fibroblasts. We identified increased expression of genes associated with fibrosis and fibrotic injury, such as *Comp, Cilp,* and *Fn1* in *Dsg2^mut/mut^*fibroblasts (22, 23), These genes were partially restored to WT levels in *Dsg2*^mut/mut^ X *Ccr2*^-/-^ mice, suggesting cross-talk between CCR2+ monocytes/macrophages and fibroblasts in the progression of myocardial fibrosis in *Dsg2^mut/mut^* mce (**Figure 6C, E**). Projection of genes differentially expressed in *Dsg2^mut/mut^* fibroblasts in the UMAP space indicated that *Postn*+ fibroblasts are a major fibroblast subset activated in the context of ACM (**Figure 6D**). These data are consistent with a known role of *Postn*+ fibroblasts in fibrosis-associated myocardial infarction and pressure overload (21, 24, 25). Pathway enrichment analysis revealed that genes upregulated in *Dsg2*^mut/mut^ fibroblasts were associated with extracellular matrix remodeling and fibrogenesis (**Figure 6E**). Consistent with our CITE-seq analysis, immunostaining showed increased *Postn* + cells in *Dsg2^mut/mut^*hearts compared to WT and *Dsg2^mut/mut^* X *Ccr2*^-/-^ hearts (**Figure 6G, H**).

**Figure 6.**
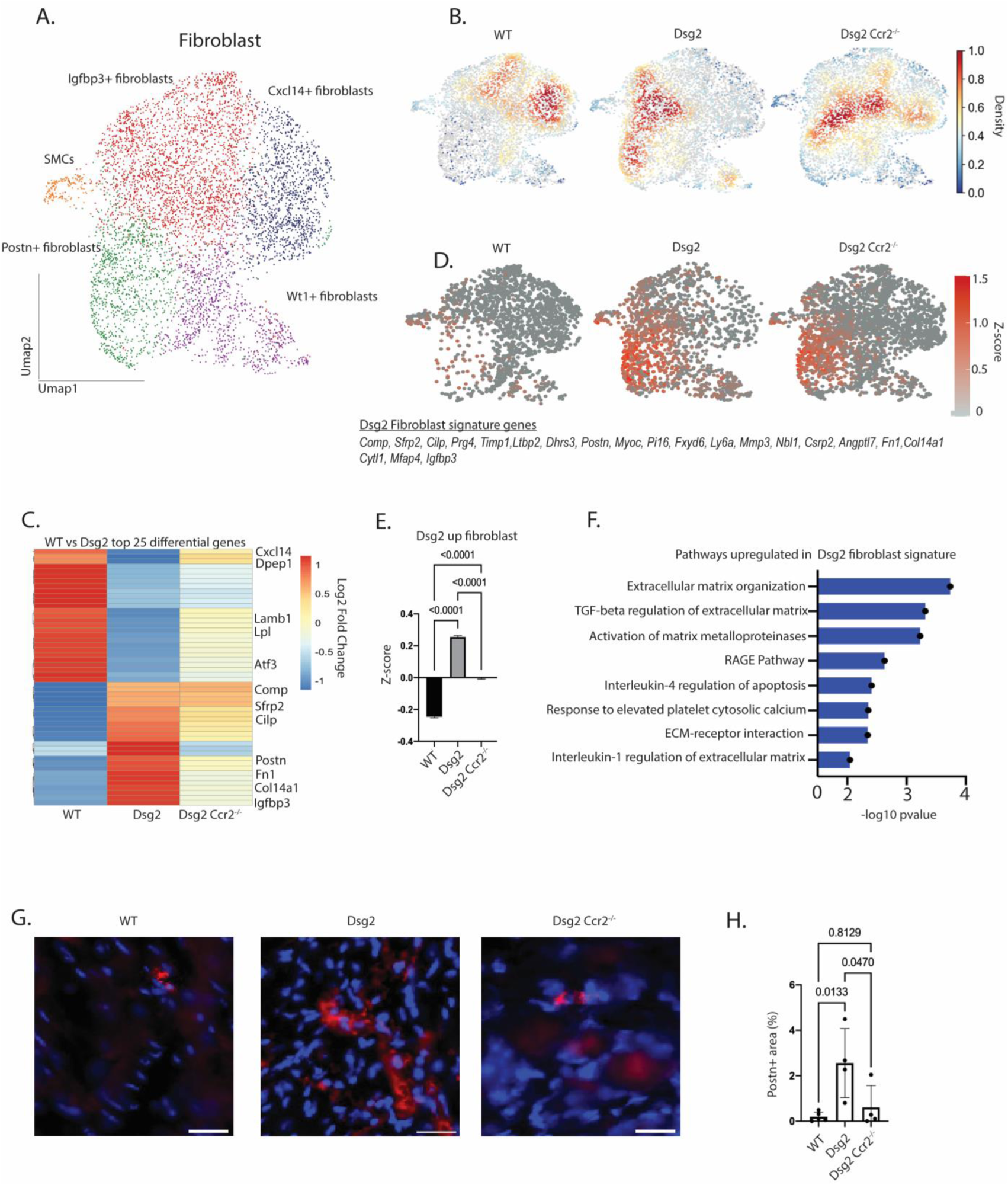
*Postn+* fibroblasts are expanded in *Dsg2*^mut/mut^ hearts through a CCR2+ monocyte- and macrophage-dependent mechanism. (**A**) Uniform Manifold Approximation and Projection (UMAP) re-clustering of fibroblast population. (**B**) Gaussian kernel density estimation of cells within the fibroblast cluster across the indicated genotypes. (**C**) Heatmap of top 25 differentially expressed genes in the fibroblast cluster between WT and *Dsg2^mut/mut^* mice with side-by-side comparison of the expression of those same genes from *Dsg2^mut/mut^* X *Ccr2^-/-^* mice. Example genes are annotated. (**D**) Z-score feature plot, overlaying a fibroblast gene signature derived from the heatmap in **C** (genes listed to the side) and displayed on the fibroblast UMAP projection, split by genotype. (**E**) Graph of z-score values for fibroblast gene signature derived from heatmap in **C** compared across genotypes (P values determined by one-way ANOVA test). (**F**) Top KEGG pathways for top 25 differentially upregulated genes in *Dsg2^mut/mut^* mice (derived from **C**). (**G**) Representative images depicting Periostin+ (red) areas via immunofluorescence staining from mice aged 16 weeks from indicated genotypes. (**H**) Quantification of Periostin+ area as a percentage of total area in mice of the indicated genotypes (P-values determined by one-way ANOVA test; WT, n=5 samples; *Dsg2^mut/mut^*, n=4 samples; *Dsg2^mut/mut^* X *Ccr2^-/-^*, n=4 samples).

### Single nucleus RNA sequencing reveals a role for CCR2+ monocytes and macrophages in cardiac myocyte remodeling in ACM

Cardiac myocytes cannot easily be isolated for single cell sequencing (15). To circumvent this limitation, we performed single nuclei RNA sequencing on pooled frozen hearts from 16-week-old WT, *Dsg2^mut/mut^* and *Dsg2^mut/mut^* X *Ccr2*^-/-^ mice (**Figure 7A**). After pre-processing and application of quality control filters (**Supplemental Figure 4A-C**) (15) we identified 10 distinct cell types (**Figure 7B, Supplemental Figure 4D, E**), including cardiac myocytes, fibroblasts, endothelial cells, pericytes/smooth muscle cells, epicardial cells, macrophages, T-cells and glia-like cells. Differential gene expression analysis revealed cell type-specific transcriptional differences between all major cell types in both WT vs *Dsg2^mut/mut^* hearts and *Dsg2^mut/mut^* vs *Dsg2^mut/mut^* X *Ccr2*^-/-^ hearts (**Supplemental Figure 4F, G**). Further analysis of the cardiac myocyte population identified three distinct clusters referred to as CM1 (“healthy” cardiac myocytes), CM2 (“failing” cardiac myocytes), and CM3 (conduction system cardiac myocytes) (15, 26) **(Figure 7D, Supplemental Figure 5A, B**). Cell composition and kernel density analysis showed that the CM2 cluster was increased in *Dsg2*^mut/mut^ hearts compared to either WT or *Dsg2^mut/mut^* X *Ccr2*^-/-^ hearts (**Figure 7C, E**). Differential gene analysis demonstrated an increase in genes associated with heart failure and inflammation including *Ankrd1, Xirp2* and *Tlr4* (15) in *Dsg2^mut/mut^* cardiac myocytes. Many of these genes were normalized to WT levels in *Dsg2^mut/mut^*X *Ccr2*^-/-^ cardiac myocytes (**Figure 7F-I**). These findings highlight cross-talk between CCR2+ monocytes/macrophages and cardiac myocytes in the pathogenesis of ACM.

**Figure 7.**
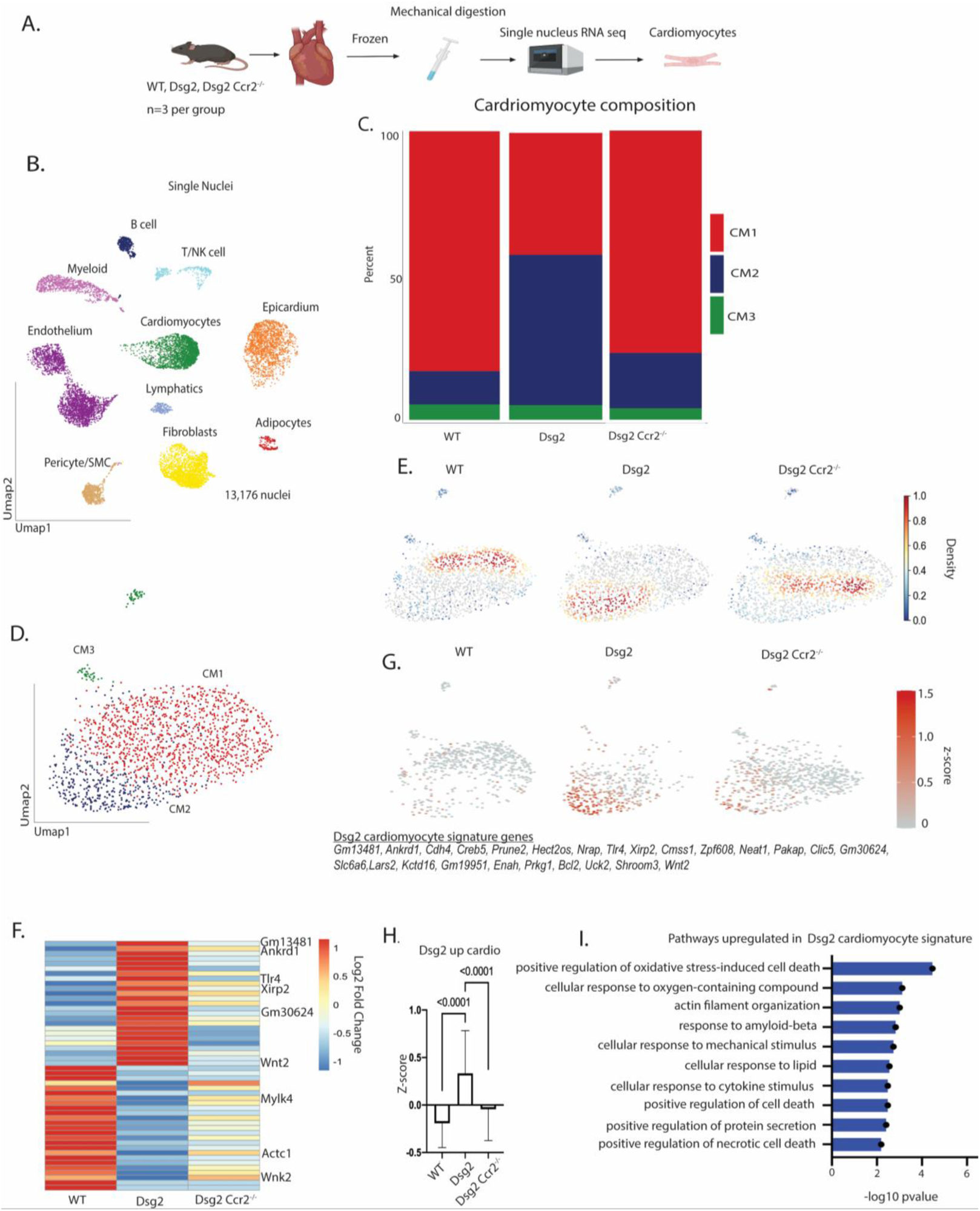
Single nucleus RNA sequencing reveals a role for CCR2+ monocytes and macrophages in cardiac myocyte remodeling in ACM. (**A**) Schematic depicting design of snRNA-seq experiments. Frozen whole hearts were mechanically homogenized. (**B**) Uniform Manifold Approximation and Projection (UMAP) of 13,176 nuclei after QC and data filtering using standard Seurat pipeline. (**C**) Composition graph showing proportion of different populations within the cardiac myocyte cluster. (**D**) Uniform Manifold Approximation and Projection (UMAP) re-clustering of cardiac myocyte population. (**E**) Gaussian kernel density estimation of cells within the cardiac myocyte cluster across the indicated genotypes. (**F**) Heatmap of top 25 differentially expressed genes in the cardiac myocyte cluster between WT and *Dsg2^mut/mut^* mice with side-by-side comparison of the expression of those same genes from *Dsg2^mut/mut^* X *Ccr2^-/-^*. Example genes are annotated. (**G**) Z-score feature plot, overlaying a cardiac myocyte gene signature derived from the heatmap in **F** (genes listed to the side) and displayed on the cardiac myocyte UMAP projection split by genotype. (**H**) Graph of z-score values for cardiac myocyte gene signature derived from heatmap in **F** compared across genotypes (P values determined by one-way ANOVA test). (**I**) Top KEGG pathways for top 25 differentially upregulated genes in *Dsg2^mut/mut^*mice (derived from **F**).

### Mechanisms of contractile dysfunction in ACM

Results in **Figure 3** showed that 16-week-old *Dsg2^mut/mut^*x *Ccr2*^-/-^ mice exhibited persistent contractile dysfunction despite having markedly reduced fibrosis and PVC burden. In contrast, contractile function was largely preserved in 16-week-old *Dsg2^mut/mut^* X DN-*IĸBα*^+/-^ mice, which also showed reduced myocardial fibrosis and PVC burden (**Figure 1**). Taken together, these results suggest that contractile dysfunction in ACM is primarily impacted by cardiac myocyte NFκB signaling. However, CCR2+ monocyte and macrophage recruitment may also contribute by promoting myocardial injury and fibrosis. To further evaluate causes of contractile dysfunction in ACM, we treated 16-week-old *Dsg2^mut/mut^* and *Dsg2^mut/mut^* X *Ccr2^-/-^* mice with Bay 11-7082, a potent inhibitor of NFκB (6), for 8 weeks and assessed contractile function and arrhythmias before and after treatment. Prior to Bay 11-7082 treatment, *Dsg2^mut/mut^* and *Dsg2^mut/mut^* X *Ccr2^-/-^*mice both showed significant and roughly equivalent reductions in LV ejection fractions (**Figure 8**). Vehicle-treated *Dsg2^mut/mut^* mice showed further deterioration of LV function during the 8-week treatment interval, whereas Bay 11-7082-treated *Dsg2*^mut/mut^ mice showed no further disease progression and actually exhibited modest improvement in LV ejection fraction. Strikingly, LV ejection fractions in Bay 11-7082 treated *Dsg2^mut/mut^* X *Ccr2^-/-^* mice were normalized to levels typically seen in WT mice (**Figure 8**). These observations indicate that LV systolic dysfunction in *Dsg2^mut/mut^* mice is caused both by negative inotropic effects of NFκB signaling in viable cardiac myocytes and myocardial injury mediated by CCR2+ monocyte and macrophage infiltration. These results highlight contributions of cardiac myocyte-intrinsic and -extrinsic mechanisms of contractile dysfunction in ACM. They also have important implications for ACM patients as they suggest that anti-inflammatory therapy may be beneficial in patients with established disease.

**Figure 8.**
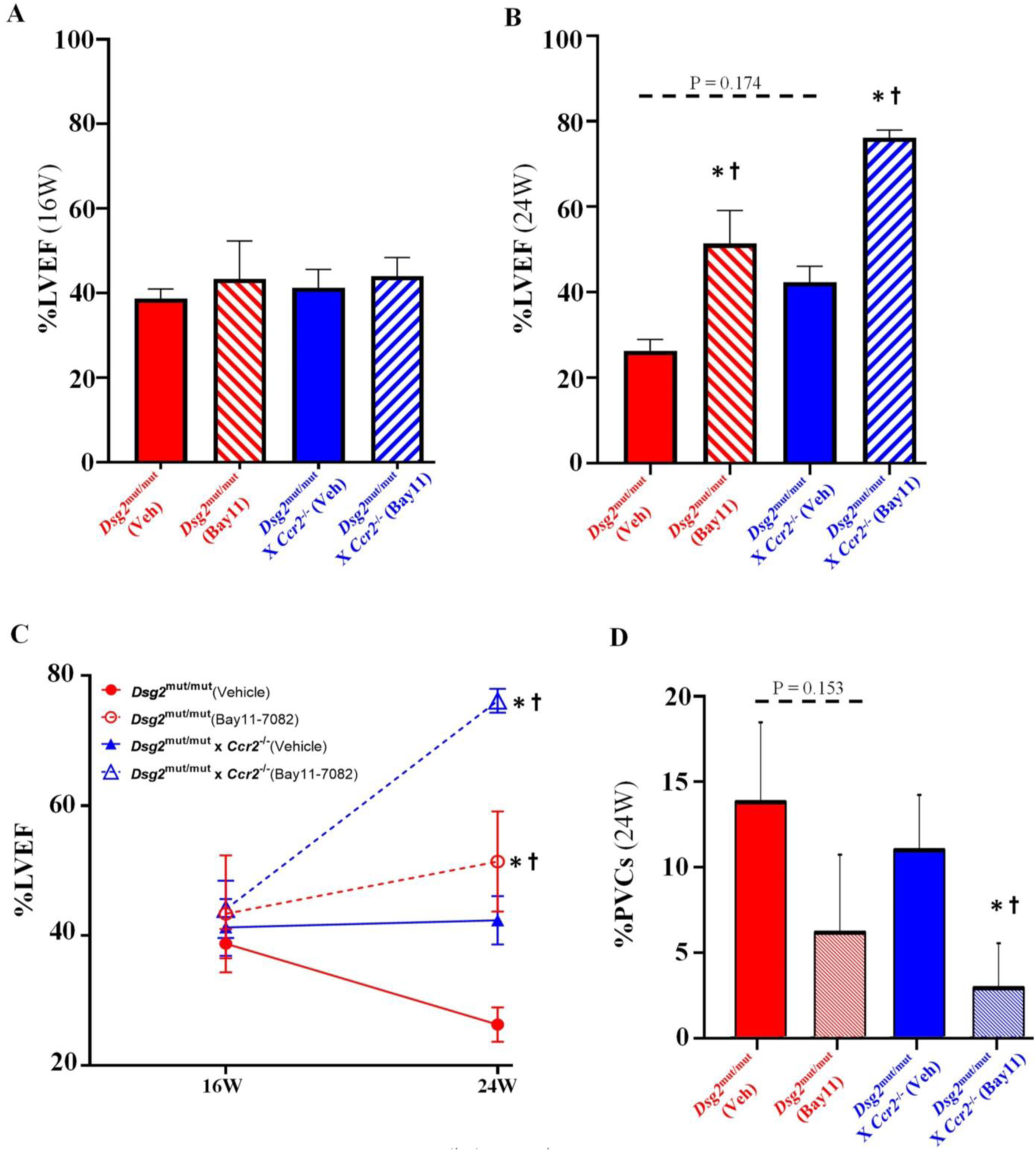
Contractile dysfunction is rescued in *Dsg2*^mut/mut^ X *Ccr2*^-/-^. (**A, B**) Percent left ventricular ejection fraction (%LVEF) at 16 (16W) and 24 weeks (24) of age in *Dsg2*^mut/mut^ and *Dsg2^mut/mut^* X *Ccr2*^-/-^ mice treated for 8 weeks with either Vehicle (Veh) or the NFкB inhibitor, Bay11-7082 (Bay11). (**C**) Longitudinal analysis of changes in %LVEF from 16 to 24 weeks. Note preserved cardiac function in Veh-treated *Dsg2^mut/mut^* X *Ccr2*^-/-^ mice, whereas Bay11-treated *Dsg2^mut/mut^* X *Ccr2*^-/-^ mice showed significant improvement in function with 8 weeks of Bay11 treatment. (**D**) Percent PVCs (%PVCs) at 24 weeks of age. *P<0.05 any cohort vs *Dsg2^mut/mut^* (Vehicle); ^†^P<0.05 Bay11-7082 treated mice vs Vehicle-treated mice within genotype; using one-way ANOVA with Tukey’s posthoc analysis.

## DISCUSSION

Inflammation has been recognized in ACM for as long as the disease has been known (4, 5). Inflammatory infiltrates are seen at autopsy in the hearts of most ACM patients, and are especially common in ACM patients who died suddenly (3, 5). They typically occur in both ventricles even if macroscopic disease is confined to the RV, and their presence may lead to the misdiagnosis of myocarditis (27, 28). A histologic picture reminiscent of acute myocarditis may reflect an active phase of ACM associated with accelerated disease progression (28). Yet, while inflammatory cells likely contribute to the pathogenesis of ACM, no previous studies have validated such a mechanism nor have specific types of inflammatory cells been implicated in myocardial injury and arrhythmias in ACM. Furthermore, iPSC-cardiac myocytes expressing ACM disease variants in *PKP2* (6) or *DSG2* (10) are known to produce large amounts of pro-inflammatory mediators under the control of NFκB, including primordial cytokines of the innate immune response such as IL-1β, INFγ and TNFα. These cytokines are expressed under basal conditions *in vitro* without stimulation or exogenous provocation and in the absence of inflammatory cells.

These observations raise the question about the relative roles of innate immune responses in cardiac myocytes vs. actions of inflammatory cells in the pathogenesis of ACM. To answer this question, we studied a well-characterized mouse model of ACM (*Dsg2^mut/mut^* mice) that exhibits progressive myocardial injury (loss of heart muscle and replacement by fibrosis), contractile dysfunction and arrhythmias. Using a genetic approach, we defined phenotypes in *Dsg2^mut/mut^*mice in which either NFκB signaling in cardiac myocytes was prevented or actions of monocytes and macrophages expressing CCR2 were blocked. We discovered complex immune mechanisms in the pathogenesis of ACM involving cross-talk between cardiac myocytes, CCR2+ macrophages and fibroblasts.

The major new insight to come from this work is that NFκB signaling in cardiac myocytes is fundamental in the pathogenesis of ACM. Blocking this pathway in cardiac myocytes alone significantly attenuated the ACM disease phenotype in *Dsg2^mut/mut^* mice. These mice showed little if any myocardial fibrosis, a marked reduction in arrhythmias and ECG abnormalities, preservation of contractile function, and greatly reduced myocardial levels of pro-inflammatory cytokines. These results indicate that NFκB signaling in cardiac myocytes drives myocardial injury, promotes arrhythmias and stimulates production of pro-inflammatory mediators.

NFκB signaling in cardiac myocytes also had a profound impact on populations of myocardial inflammatory cells and fibroblasts in *Dsg2^mut/mut^* mice. Hearts of these mice contained reduced numbers of Lyve1+ cardiac resident macrophages, and greatly increased numbers of pro-inflammatory CCR2+ macrophages and pro-fibrotic *Postn*+ fibroblasts compared to WT mice. This highly pro-inflammatory cellular landscape was greatly attenuated when NFκB signaling in cardiac myocytes was blocked in *Dsg2^mut/mut^* X DN-*IĸBα*^+/-^ mice. These findings strongly suggest that signals emanating from cardiac myocytes in *Dsg2^mut/mut^* mice cause CCR2+ macrophages to accumulate in the heart and promote development of a subset of fibroblasts known to participate in myocardial fibrosis.

To gain insights into the actions of CCR2+ cells, we analyzed phenotypes in *Dsg2^mut/mut^* mice with germline deletion of *Ccr2.* As seen in *Dsg2^mut/mut^* X DN-*IĸBα*^+/-^ mice in which NFκB signaling in cardiac myocytes was blocked, *Dsg2^mut/mut^* X *Ccr2^-/-^* mice exhibited little if any myocardial fibrosis. They also contained fewer *Postn* + cells in their hearts than *Dsg2^mut/mut^* mice and showed significant reductions in arrhythmias and cytokine levels. These observations indicate that CCR2+ cells mediate myocardial injury and promote arrhythmias, as these disease features were clearly mitigated in *Dsg2^mut/mut^* X *Ccr2^-/-^* mice. They also suggest that the greatly reduced myocardial injury seen in *Dsg2^mut/mut^* X DN-*IĸBα*^+/-^ mice was due mainly, if not entirely, to diminished numbers of CCR2+ macrophages in these hearts. Thus, NFκB signaling in cardiac myocytes stimulates CCR2+ cells to accumulate in the heart where they mediate myocardial injury and promote arrhythmias.

Despite the distinct lack of myocardial fibrosis in *Dsg2^mut/mut^* X *Ccr2^-/-^* mice, LV contractile function was reduced to an extent seen in *Dsg2^mut/mut^*mice. This observation suggested that NFκB signaling in cardiac myocytes, which was presumably unaffected in *Dsg2^mut/mut^* X *Ccr2^-/-^* mice, caused contractile dysfunction related to negative inotropic effects of inflammation in viable myocardium. Such a mechanism was supported by studies showing that treating 16-week-old *Dsg2^mut/mut^* X *Ccr2^-/-^* mice with the NFκB blocker, Bay11-7082, fully normalized contractile function. These observations suggest that contractile dysfunction in ACM patients is related not only to loss of myocardium and its replacement by fibrofatty scar tissue, but to potentially reversible changes in viable myocardium caused by persistent innate immune signaling. If so, then anti-inflammatory therapy in ACM patients with established disease might result in at least some recovery of contractile function.

Results of CITE-seq and snRNA-seq studies revealed additional insights highlighting remarkable cross-talk between cardiac myocytes, CCR2+ macrophages and *Postn*+ fibroblasts. For example, *Tlr4*, the gene for the major pattern recognition receptor on cardiac myocytes, was one of the most highly upregulated genes in cardiac myocytes in *Dsg2^mut/mut^* mice, consistent with the chronic, non-resolving NFκB signaling known to occur in cardiac myocytes in ACM. Yet, *Tlr4* expression was not increased in cardiac myocytes in *Dsg2^mut/mut^* X *Ccr2^-/-^* mice. Similarly, *Postn* expression was greatly increased in fibroblasts in *Dsg2^mut/mut^* mice but not in *Dsg2^mut/mut^* X *Ccr2^-/-^* mice. These results suggest that signals from CCR2+ cells directly or indirectly regulate gene expression in cardiac myocytes and fibroblasts in *Dsg2^mut/mut^* mice.

The results of this study reveal new insights into mechanisms of immune injury in ACM and demonstrate bidirectional interactions between cardiac myocytes, inflammatory cells and fibroblasts. They also identify potential new drug targets and strategies. However, as novel and provocative as these findings may be, major questions remain unanswered and may serve as priorities for future research. For example, how variants in desmosomal genes lead to persistent activation of innate immune responses that fail to resolve is unknown. GSK3β, which is aberrantly activated in cardiac myocytes in ACM due to altered Wnt/β-catenin pathways (7, 29, 30), could be a mechanistic link. Activation of GSK3β has been shown to promote inflammation through NFκB, whereas inhibition of GSK3β limits inflammation (31–33). Another unanswered question concerns the pathogenic roles of specific inflammatory mediators produced by cardiac myocyte and/or inflammatory cells in ACM. Progress in this area could impact future drug discovery. Related unanswered questions concern the specific signals used by cardiac myocytes to mobilize inflammatory cells and orchestrate their injurious activities in ACM, and mechanisms used by CCR2+ cells that regulate gene expression in cardiac myocytes and fibroblasts. Finally, it should be emphasized that these studies focused only on the role of CCR2+ cells in *Dsg2^mut/mut^* mice. Other types of inflammatory cells undoubtedly participate in the pathogenesis of ACM as well.

Studies here were performed entirely in genetically manipulated mouse models of ACM. Previous studies of ACM patient-derived iPSC-cardiac myocytes suggest that inflammatory signaling mediated by NFκB is activated in a cell-autonomous fashion in ACM (6, 10). To determine if NFκB is activated in cardiac myocytes in patients with ACM, we performed parallel studies in post-mortem hearts from ACM patients who died suddenly. As reported in a companion paper, we observed nuclear signal for RelA/p65 in cardiac myocytes in a great majority of patient hearts indicating active NFκB signaling in cardiac myocytes at the time of death. This was associated with a significant increase in the number of CCR2+ cells in ACM patient hearts compared to hearts of age-matched controls with no history of heart disease. We also found evidence of activation of NFκB signaling in buccal mucosa cells obtained from young ACM patients at the time they first exhibited clinical manifestations of disease. Taken together, new insights gained here through studies of experimental models combined with patient data provide a compelling rationale for the potential benefit of anti-inflammatory therapy in ACM.

## METHODS

### Generation of double-mutant ACM mouse models

To create ACM mice in which activation of NFĸB signaling in cardiac myocytes was prevented, we crossed *Dsg2^mut/mut^* mice with mice with transgenic cardiac myocyte-specific expression of a dominant-negative form of the NFĸB pathway protein IĸBα involving deletion of 36 NH_2_-terminal amino acids (generously supplied by Dr. Douglas Mann) (11). This deletion prevents phosphorylation of Ser32/36 phosphorylation in IĸBα and subsequent nuclear localization of NFĸB in cardiac myocytes (34, 35). This line has been used to define the role of NFκB in cardiac myocyte apoptosis following ischemia (11). Mice were crossbred to obtain a line with *Dsg2*-mutant homozygosity and cardiac myocyte-specific expression of dominant-negative IĸBα (DN-*IĸBα*^+/-^) (11), thus creating *Dsg2^mut/mut^*X DN-*IĸBα*^+/-^ double-mutant mice. To produce ACM mice in which actions of CCR2+ cells were blocked, we crossed *Dsg2^mut/mut^* mice with mice with germline deletion of murine *Ccr2 (Ccr2*^-/-^, Jackson Laboratory, strain: 017586) (36). These mice were crossbred to double homozygosity to produce *Dsg2^mut/mut^* X *Ccr2*^-/-^ double-mutant mice.

### Characterization of disease phenotypes

Phenotypes in *Dsg2^mut/mut^* and the two double-mutant lines were studied in mice at 8 and 16 weeks of age and compared to age-matched wild-type (WT) controls. As previously reported, 8-week-old *Dsg2^mut/mut^*mice exhibit only a modest disease phenotype consisting of a few premature ventricular complexes (PVCs), minimally reduced contractile function and little if any myocardial injury (6, 7). By 16 weeks of age, however, they show significant arrhythmias, marked contractile dysfunction and extensive myocardial replacement fibrosis (7). Accordingly, to define the effects of NFκB signaling in cardiac myocytes and actions of CCR2+ cells in the development of the ACM phenotype, all mice underwent echocardiographic and ECG analyses at 8 and 16 weeks of age. Then, hearts obtained from 16-week-old animals were further analyzed by: 1) histology, to measure the amount of ventricular fibrosis; 2) immunohistochemistry and RNA-scope *in situ* hybridization, to measure the number of specific subsets of macrophages; 3) cytokine arrays, to measure the levels of various inflammatory mediators; and 4) single nuclei RNA sequencing (sn-RNAseq) and cellular indexing of transcriptomes and epitomes (CITE-seq) to characterize gene expression and related changes in specific cells types in the heart including macrophages, fibroblasts and cardiac myocytes. Detailed descriptions of these methods are included in supplemental materials.

### In vivo drug treatment

The effects of Bay 11-7082, a small molecule inhibitor of NFĸB, on left ventricular contractile function were compared in *Dsg2^mut/mut^* mice and *Dsg2^mut/mut^* X *Ccr2*^-/-^ mice to determine the extent to which inhibition of NFĸB signaling could prevent further disease progression and promote recovery of cardiac function. 16-week-old *Dsg2^mut/mut^*and *Dsg2^mut/mut^* X *Ccr2*^-/-^mice underwent echocardiography before being implanted with subcutaneous osmotic minipumps (Alzet, Model 1004), as previously described (6). They received either vehicle or drug (50 µg/µL Bay11-7082 dissolved in 65% dimethyl sulfoxide, 15% ethanol, and 20PBS). Drug-treated mice received 5 mg/kg/day Bay11-7082 via continuous infusion (0.11 µL/hr for 28 days); vehicle-treated mice received an equivalent volume of vehicle for 28 days. Minipumps were replaced at 20 weeks of age, and treatment was continued for an additional 4 weeks. Final echocardiograms were obtained in both groups at 24 weeks of age.

### Statistical analysis

All data are presented as mean±SEM; n-values and the statistical analyses performed for each experiment are indicated in figure legends and tables. Gaussian distributions were assumed considering sample sizes (n≥10). Differences in measured variables were assessed with Mann-Whitney or 1-way ANOVA with Tukey post-hoc analysis. A P-value of <0.05 was considered statistically significant. All statistical analyses were analyzed using GraphPad Prism (v9.2) software.

### Study approval

All experiments in this study conformed to the National Institutes of Health Guide for the Care and Use of Laboratory Animals (NIH publication no. 85-23, revised 1996). All protocols were approved by the Animal Care and Use Committees at Florida State University and Washington University in St. Louis. All animals were housed in a 12-hour-light/dark cycle, climate-controlled facility with *ad libitum* access to water and standard rodent chow.

### Data Availability

Raw sequence files that support the findings of this study are available on the Gene Expression Omnibus (GSE228048). Processed data are available upon request from the authors.

## ACKNOWLEDGMENTS

We thank Dr. Douglas Mann (Washington University School of Medicine in St. Louis) from whom we acquired the DN-*IĸBα*^+/-^ mice. This work was supported by an American Heart Association Career Development Award (19CDA34760185, S.P. Chelko); a National Institutes of Health grant (R01-HL148348, J.E. Saffitz); a Washington University in St. Louis Rheumatic Diseases Research Resource-Based Center grant (NIH P30AR073752, K. Lavine); a National Institutes of Health grant (R35 HL161185, K. Lavine); a Leducq Foundation Network grant (#20CVD02, K. Lavine); a Burroughs Welcome Fund grant (1014782, K. Lavine); a Children’s Discovery Institute of Washington University and St. Louis Children’s Hospital grant (CH-II-2015-462, CH-II-2017-628, PM-LI-2019-829, K. Lavine); a Foundation of Barnes-Jewish Hospital grant (8038-88, K. Lavine); and generous gifts from Washington University School of Medicine (K. Lavine). V. Penna is supported by a National Institutes of Health grant (5T32AI007163-44, V. Penna).

## SUPPLEMENTAL MATERIALS: FIGURES

**Supplemental Figure 1. Quality Control and Overview of CITE-seq data**

**(A)** Violin plot of the RNA count per cell, split by genotype. **(B)** Violin plot of the number of genes per cell, split by genotype. **(C),** Violin plot of the percent of mitochondrial reads per cell split by genotype. **(D)** Violin plot displaying characteristic marker gene for each identified cell population. **(E)** Composition graph showing proportion of different populations within the entire CITE-seq data set. **(F)** Dot plot showing differentially expressed genes in each cell population between WT and *Dsg2^mut/mut^* mice. **(G)** Dot plot showing differentially expressed genes in each cell population between *Dsg2^mut/mut^* and *Dsg2^mut/mut^* X *Ccr2^-/-^*. **(H)** Gaussian kernel density estimation of cells within the entire CITE-seq data set across the indicated genotypes.

**Supplemental Figure 2. Characterization of Macrophage/monocyte clusters**

**(A)** Heat map of the top 10 genes by log 2-fold change enriched in each cluster. **(B)** Z-score feature plots for transcriptional signatures enriched in each sub cluster within the macrophage/monocyte population. Genes used for identification were selected based on enrichment from Seurat differential expression analysis.

**Supplemental Figure 3. Characterization of Fibroblast clusters**

**(A)** Heat map of the top 10 genes by log 2-fold change enriched in each cluster. **(B)** Composition graph showing proportion of different populations within the fibroblast cluster. **(C)** Z-score feature plots for transcriptional signatures enriched in each sub-cluster within the fibroblast population. Genes used for identification were selected based on enrichment from Seurat differential expression analysis.

**Supplemental Figure 4. Quality Control and Overview of snRNA-seq data**

**(A)** Violin plot of the RNA count per cell, split by genotype. **(B)** Violin plot of the number of genes per cell, split by genotype. **(C)** Violin plot of the percent of mitochondrial reads per cell split by genotype. **(D)** Violin plot displaying characteristic marker gene for each identified cell population. **(E)** Composition graph showing proportion of different populations within the entire snRNA-seq data set. **(F)** Dot plot showing differentially expressed genes in each cell population between WT and *Dsg2^mut/mut^*mice. **(G)** Dot plot showing differentially expressed genes in each cell population between *Dsg2^mut/mut^* and *Dsg2^mut/mut^* X *Ccr2^-/-^* mice. **(H)** Gaussian kernel density estimation of cells within the entire snRNA-seq data set across the indicated genotypes.

**Supplemental Figure 5. Characterization of Cardiac myocyte clusters**

**(A)** Heat map of the top 10 genes by log 2-fold change enriched in each cluster. **(B)** Z-score feature plots for transcriptional signatures enriched in each sub cluster within the cardiac myocyte population. Genes used for identification were selected based on enrichment from Seurat differential expression analysis.

**Supplemental Figure 6. Time course of disease in *Dsg2^mut/mut^* mice and NFκB inhibition study design.** PD1, postnatal day 1; W, week; ↓%EF, reduced percent ejection fraction; mouse clip art, Alzet minipump implantation at 16W and 20W of age.

**Supplemental Table 1: Fold-changes in cytokine levels in the hearts of *Dsg2^mut/mut^*, *Dsg2^mut/mut^* X DN-*IĸBα^+/-^*, and *Dsg2^mut/mut^* X *Ccr2^-/-^* mice.** Data are presented as fold-change and P-values as compared to age-matched WT mice (n=5 mice/cohort). P-values were determined using one-way ANOVA with Tukey’s posthoc test.

## SUPPLEMENTAL MATERIALS: METHODS

**Echocardiography** was performed using a 2100 Vevo Visualsonic machine equipped with a 40 MHz ultrahigh frequency linear array microscan transducer. Images were obtained according to the American Society of Echocardiography guidelines for animals (37). Short-axis, m-mode and parasternal long-axis images, B-mode images at the level of the papillary muscles were acquired at a sweep speed of 200 mm/s, as previously described (6–8). Measurements obtained from three to five echocardiographic images for each mouse were averaged to assess left ventricular ejection fraction (EF) and wall and chamber dimensions (6–8).

**Body surface ECGs** were performed in lightly anesthetized mice (nose cone delivery of 1.5 - 2% isoflurane vaporized in 100% O_2_) using PowerLab to obtain Lead I ECG recordings, as previously described (6–8). Recordings were analyzed using the ECG Analysis Add-on Software from LabChart Pro (LabChart Pro 8, MLS360/8, AD Instruments). Signal-averaged ECGs (SAECGs) constructed from 10-minute recordings using LabChart were used to measure ECG durations, intervals, and wave and amplitude parameters. The entire 10-minute ECG recording for each animal was analyzed to determine the percentage of PVCs by total beats. Once 16-week ECG studies were completed, mice were euthanized and hearts excised for histologic, immunohistochemical, cytokine and sequencing analyses.

**Myocardial fibrosis** was measured in long-axis sections of formalin-fixed, paraffin-embedded hearts cut at 5µm and stained with Masson’s trichrome stain. The total amount of right and left ventricular section area occupied by fibrosis was determined by the sum of all fibrotic areas divided by total myocardial area using ImageJ version 1.53e software (8).

**Immunohistochemistry** was performed to measure the number of CD68 and Lyve1-expressing (15, 38) cells in hearts of WT, *Dsg2^mut/mut^* and double mutant mice at 16 weeks of age. Isolated hearts were perfused with 1X PBS and fixed overnight in 4% paraformaldehyde in 1X PBS at 4°C. After being dehydrated in 30% sucrose in PBS at 4°C overnight, they were embedded in O.C.T. (Sakura, Cat. No. 4583) and stored at -80⁰C for 10 min. They were then sectioned at 12 µm using a Leica cryostat, mounted on positively charged slides, and stored at -20⁰C. For staining, the following steps were performed at room temperature and protected from light. Slides were brought to room temperature for 5 min and washed in 1X PBS for 5 min. Sections were permeabilized in 0.25% Triton-X in 1X PBS for 5 min and blocked in 5% BSA in 1X PBS for 1 hr. Sections were then stained with primary antibodies (rat anti-CD68, BioLegend Cat. No. 137002, at 1:400; rabbit anti-Lyve1, Abcam Cat. No. 14917, at 1:200; or rabbit anti-Periostin, Abcam Cat. No. 215199, at 1:200) diluted in 1% BSA in 1X PBS for 1 hr. Sections were then washed 3 times for 5 min each with 1X PBS-Tween. After washing, appropriate secondary antibodies (goat anti-rat Alexa Fluor 555, Invitrogen Cat. No. A21434; donkey anti-rabbit Alexa Fluor 647, Abcam Cat. No. ab150075; or goat anti-rabbit Alexa Fluor 555, Invitrogen Cat. No. A21428) were added for 1 hr at 1:1000 in 1% BSA in 1X PBS. Slides were again washed 3 times for 5 min each with 1X PBS-T, and then mounted with DAPI mounting media (Millipore Sigma, Cat. No. F6057) and cover slipped. Slides were stored at 4⁰C, protected from light, and imaged within 1 week using the 20x objective lens of a Zeiss LSM 700 confocal microscope. For quantification, ten 20x fields were prepared from each mouse heart. Cell numbers and area percentages were quantified in Zen Blue, and data were displayed as the average per 20x field per mouse.

**RNA-scope *in situ* hybridization** was performed as previously (15), which was used to identify and quantify CCR2+ cells in hearts of WT, *Dsg2*^mut/mut^ and double mutant mice at 16 weeks of age. Hearts were fixed and dehydrated, and sections were cut as above. CCR2 (Cat. No. 433271) RNA-Scope probes produced by ACDBio were used. RNA was visualized using RNA-Scope Multiplex Fluorescent Reagent kit v2 Assay (ACDBio 323100). Fluorescent images were collected using a Zeiss LSM 700 laser scanning confocal microscope and quantification was performed as above.

**Myocardial cytokines** were measured using Proteome Profiler Mouse XL Cytokine Array Kits (R&D Systems, Cat. No. ARY028) (6). Frozen heart samples were lysed in RIPA buffer containing 1:100 protease and phosphatase inhibitor cocktails. Protein content in lysates were quantified via BCA assay and 40µg/µl of protein lysate were probed on cytokine array blots. Following the manufacturer’s protocol, blots were incubated with ECL substrate, imaged on an Azure Biosystems 400 imager and analyzed using Quick Spots image analysis software (Version 25.5.1.2, Ideal Eyes Systems).

**Sample preparation for CITE-seq** was performed as previously (15, 21). Freshly isolated hearts from 16-week-old WT, *Dsg2*^mut/mut^ and *Dsg2^mut/mut^* X *Ccr2*^-/-^ mice (n=3 per group) were minced on ice with a razor blade, transferred to a 15mL conical tube containing 3mL DMEM with 170µL collagenase IV (250U/mL final concentration), 35µL DNAse1 (60U/mL), and 75µL hyaluronidase (60U/mL), and incubated at 37°C for 45 min with agitation. Thereafter, the digestion reaction was quenched with 5mL of HBB buffer (2% FBS and 0.2% BSA in HBSS), passed through 40µm filters into a 50mL conical tube and transferred back into a 15mL conical tube to obtain tighter pellets. Samples were then spun down at 4°C, 1200 rpm for 5 min and the supernatant was discarded. Pellets were resuspended in 1mL ACK Lysis buffer (Gibco, Cat. No. A10492-01) and incubated at room temperature for 5 min, followed by the addition of 9mL DMEM and centrifugation (4°C, 5 min, 1200 rpm). Supernatant was discarded and the pellets were resuspended in 2mL FACS buffer (2% FBS and 2mM EDTA in calcium/magnesium free PBS); centrifugation was repeated in above conditions and supernatant aspirated. The TotalSeq A 277 panel (BioLegend, Cat. No. 199901) antibody cocktail was resuspended in 100µL of FACS buffer and added to each sample. The combined 100µL were used to resuspend the pellet with the addition of 1µL of DRAQ5 (5mM solution Thermo Fisher Scientific, Cat. No. 564907) and incubated on ice for 30 min. Solution was washed 3x with FACS buffer following same centrifugation as above and then resuspended in 300µL of FACS buffer and 1uL DAPI (BD Biosciences, Cat. No. 564907) and filtered into filter top FACS tubes. First singlets were gated and subsequent DRAQ5+/DAPI-events were collected in 300µL cell resuspension buffer (0.04% BSA in PBS) – collected cells were centrifuged as above and resuspended in collection buffer to a target concentration of 1,000 cells/µL. Cells were counted on a hemocytometer before proceeding with the 10x protocol.

**Sample preparation for single nuclei RNA-seq (snRNA-seq):** was performed as previously (15), using frozen hearts from 16-week-old WT, Dsg2^mut/mut^ and *Dsg2^mut/mut^* X *Ccr2*^-/-^ mice (n=3 per group) were minced with a razor blade and transferred into a 5mL Dounce homogenizer containing 1– 2mL chilled lysis buffer (10 mM Tris-HCl, pH 7.4, 10 mM NaCl, 3 mM MgCl_2_ and 0.1% NP-40 in nuclease-free water). Samples were gently homogenized using five passes without rotation, and then incubated on ice for 15 min. Lysate was gently passed through a 40μm filter into 50mL conical tube, followed by rinsing the filter once with 1ml lysis buffer and transfer of lysate to a new 15mL conical tube. Nuclei were then centrifuged at 500*g* for 5 min at 4°C, followed by resuspension in 1mL nuclei wash buffer (2% BSA and 0.2 U /µL RNase inhibitor in 1X PBS) and filtered through a 20μm pluristrainer into a fresh 15mL conical tube. Centrifugation was repeated according to the above parameters. Supernatant was then removed, and nuclei were resuspended in 300µL nuclei wash buffer and transferred to a 5mL tube for flow sorting. Then, 1µl DRAQ5 (5 mM solution; Thermo Fisher, Cat. No. 62251) was added, mixed gently and allowed to incubate for 5 min before sorting. DRAQ5^+^ nuclei were sorted into nuclei wash buffer. Recovered nuclei were centrifuged again under the above parameters and were gently resuspended in nuclei wash buffer to a target concentration of 1,000 nuclei/µL. Nuclei were counted on a hemocytometer.

**CITE-seq library preparation:** was performed as previously (15, 21). Collected cells were processed using the single Cell 3’ Kit v 3.1 (10x Genomics PN: 1000268). 10,000 cells were loaded onto ChipG (PN: 1000121) for GEM generation. Reverse transcription, barcoding and complementary DNA amplification of the RNA and ADT tags were performed as recommended in the 3’ v3.1 chromium protocol. Single-cell libraries were prepared using the single Cell 3’ Kit v 3.1 following a modified 3’ v3.1 assay protocol (User Guide CG000206) to concurrently prepare gene expression and TotalSeq A antibody derived tag (ADT) libraries as recommended by BioLegend. 1µl of 0.2uM ADT Additive Primer (CCTTGGCACCCGAGAATT*C*C) and 15µl of cDNA Primers (PN: 2000089) were used to amplify cDNA. ADT libraries were amplified with a final concentration of 0.25 µM SI Primer (AATGATACGGCGACCACCGAGATCTACACTCTTTCCCTACACGACGC*T*C) and 0.25 µM TrueSeq Small RNA RPI primer (CAAGCAGAAGACGGCATACGAGAT [6nt index] GTGACTGGAGTTCCTTGGCACCCGAGAATTC*C*A) using 11 cycles. Gene expression libraries were indexed using Single Index Kit T Set A (PN: 2000240). Libraries were sequenced on a NovaSeq 6000 S4 flow cell (Illumina).

### snRNA-seq library preparation

Nuclei were processed using the Chromium Single Cell 5ʹ Reagent V2 kit from 10X Genomics (PN-1000263), as previously described (15). A total of 10,000 nuclei per sample were loaded into a Chip K for GEM generation. Reverse transcription, barcoding, complementary DNA amplification and purification for library preparation were performed according to the Chromium 5ʹ V2 protocol. Sequencing was performed on a NovaSeq 6000 platform (Illumina).

### CITE-seq alignment, quality control and cell type annotation

Raw fastq files were aligned to the mouse GRCh38 reference genome (v) using CellRanger (10x Genomics, v6.1) with the antibody capture tag for the TotalSeqA 277 antibodies, as previously described (15, 21). Subsequent quality control, normalization, dimensional reduction, and clustering were performed in Seurat v4.0. Following normalization, quality control was performed and cells passing the following criteria were kept for downstream processing: 200 < nFeature_RNA < 6000 and 1,000 < nCount_RNA < 30,000 and percentage mitochondrial reads < 10%. Raw RNA counts were normalized and scaled using SCTransform regressing out percent mitochondrial reads and nCount RNA. Principal component analysis was performed on normalized RNA counts and the number of PCs used for further processing was determined by fulfilling the following criteria: Principle components (PCs) exhibited cumulative percent >90% and the percent variation associated with the PCs was < 5%. Weighted nearest neighbor clustering (WNN) was performed with the significant RNA PCs and normalized proteins directly without PCA as previously outlined with the FindMultiModalNeighbors function in Seurat. Subsequently, a uniform manifold approximation (UMAP) embedding was constructed and FindClusters was used to unbiasedly cluster cells. Clustering was performed for a range of different resolutions (0.1-0.8 at 0.1 intervals) and differential gene expression using the FindAllMarkers function and a Wilcoxon Rank Sum test with a logFC cutoff of 0.25 and a min.pct cut-off of 0.1. Clusters were annotated using canonical gene and protein markers and subsequent violin plots were created to assess clean separation of clusters into distinct cell types. To cluster cell types into distinct cell states, the cell type of interest was subsetted, re-normalized, computed PCAs, computed UMAPs, and clustered data at a range of resolutions. DE analysis was then used to identify marker genes for each cell state and constructed a dot-plot or heatmap to assess clustering separation. Using the top marker genes, gene set z-scores were calculated and plotted in UMAP space.

**snRNA-seq alignment, quality control and cell type annotation** were performed as previously described (15, 21). Raw fastq files were aligned to the mouse GRCh38 reference genome (v) using CellRanger (10x Genomics, v6.1). Subsequent quality control, normalization, dimensional reduction, and clustering were performed in Seurat v4.0. Following normalization, quality control was performed and cells passing the following criteria were kept for downstream processing: 500 < nFeature_RNA < 4000 and 1,000 < nCount_RNA < 16,000 and percentage mitochondrial reads < 3%. To remove doublets, Scrublet was run with a cutoff score of >0.25 to identify doublets. Following doublet removal, raw RNA counts were normalized and scaled using SCTransform. PCs were then calculated, and an elbow plot was generated to select the cutoff for significant PCs to use for downstream analysis. UMAP dimensional reduction was then computed using the selected significant PCs. Unsupervised clustering was then performed using the FindNeighbors and FindClusters function, again using the selected significant PC level as above, calculating clustering at a range of resolutions between 0.1–0.8 at intervals of 0.1. Differential gene expression was performed using the FindAllMarkers command and a Wilcoxon Rank Sum test with a logFC cutoff of 0.25 and a min.pct cut-off of 0.1. Clusters were annotated in the same manner as the CITE-seq analysis. Subsequent sub-clustering and DE analysis was performed in the same manner as above.

### Density shift calculations

R was used to compute cell type composition across genotypes. To assess shifts in cell density within both the global object and individual cell types, the .rds object was converted to a .h5ad file format and scanpy.tl.embedding function which employs a Gaussian kernel density estimation of cell number was used within the UMAP embedding, as previously described (15, 21). Density values were scaled from 0-1 within that category.

### Pathway analysis

Statistically significant DE genes were used to perform pathway analysis via EnrichR (https://maayanlab.cloud/Enrichr/). Pathway enrichment values were downloaded as .csv files and plots generated in Prism.

