## Supplementary figures and images for "Mechanisms of Innate Immune Injury in Arrhythmogenic Cardiomyopathy"

### supp3.tiff

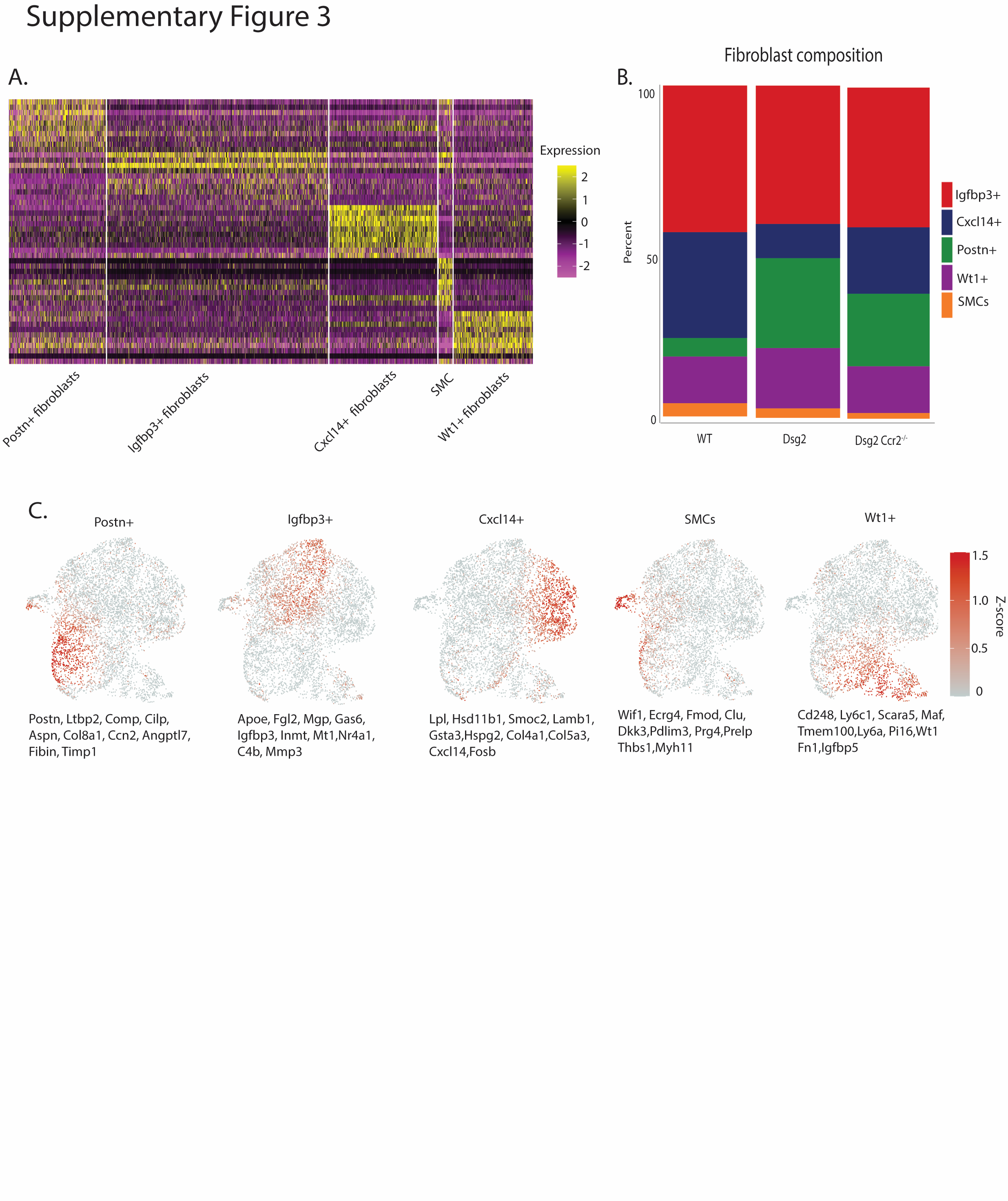

### supp4.tiff

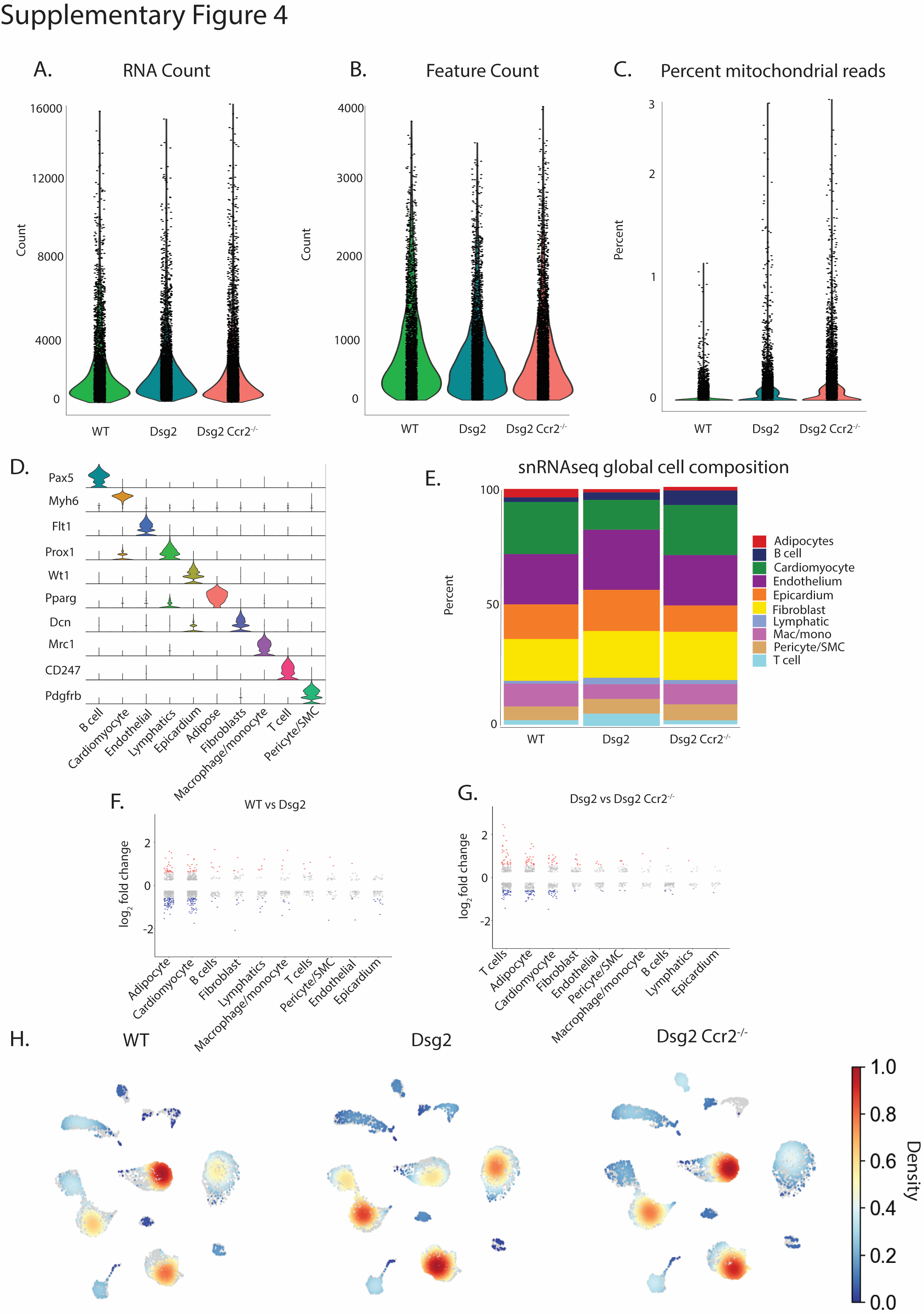

### supp5.tiff

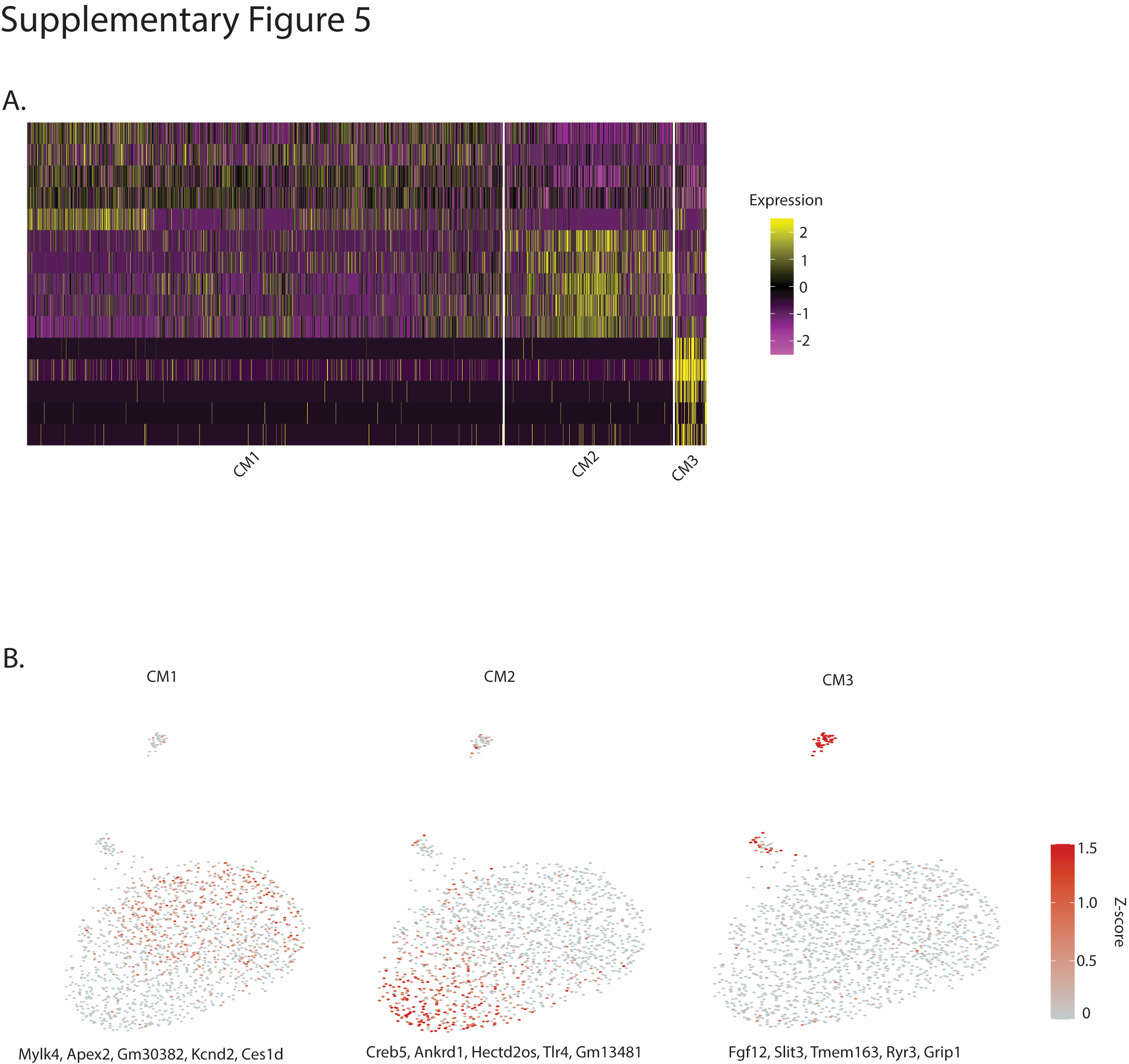

### supp-1.tiff

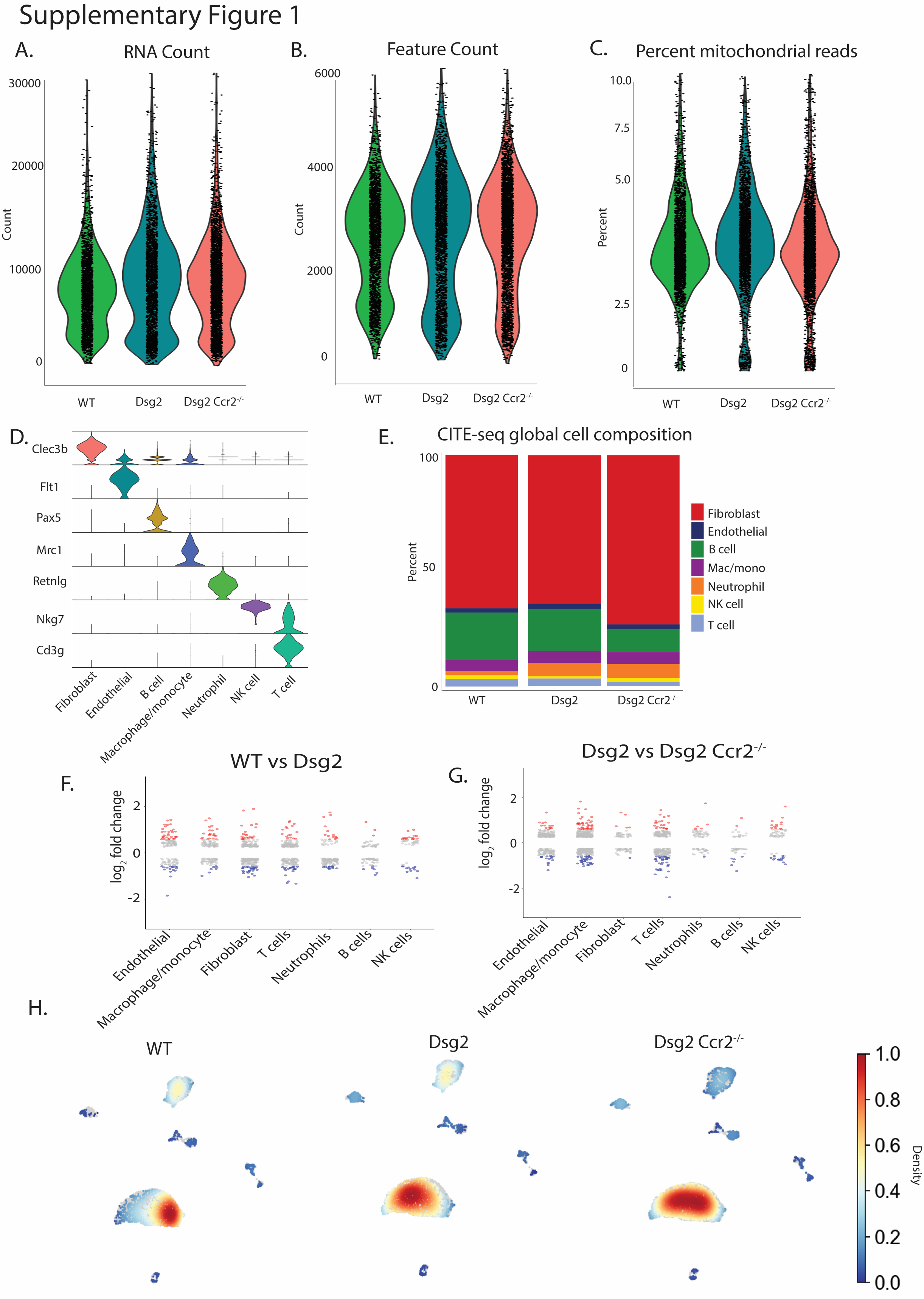

### supp-2.tiff

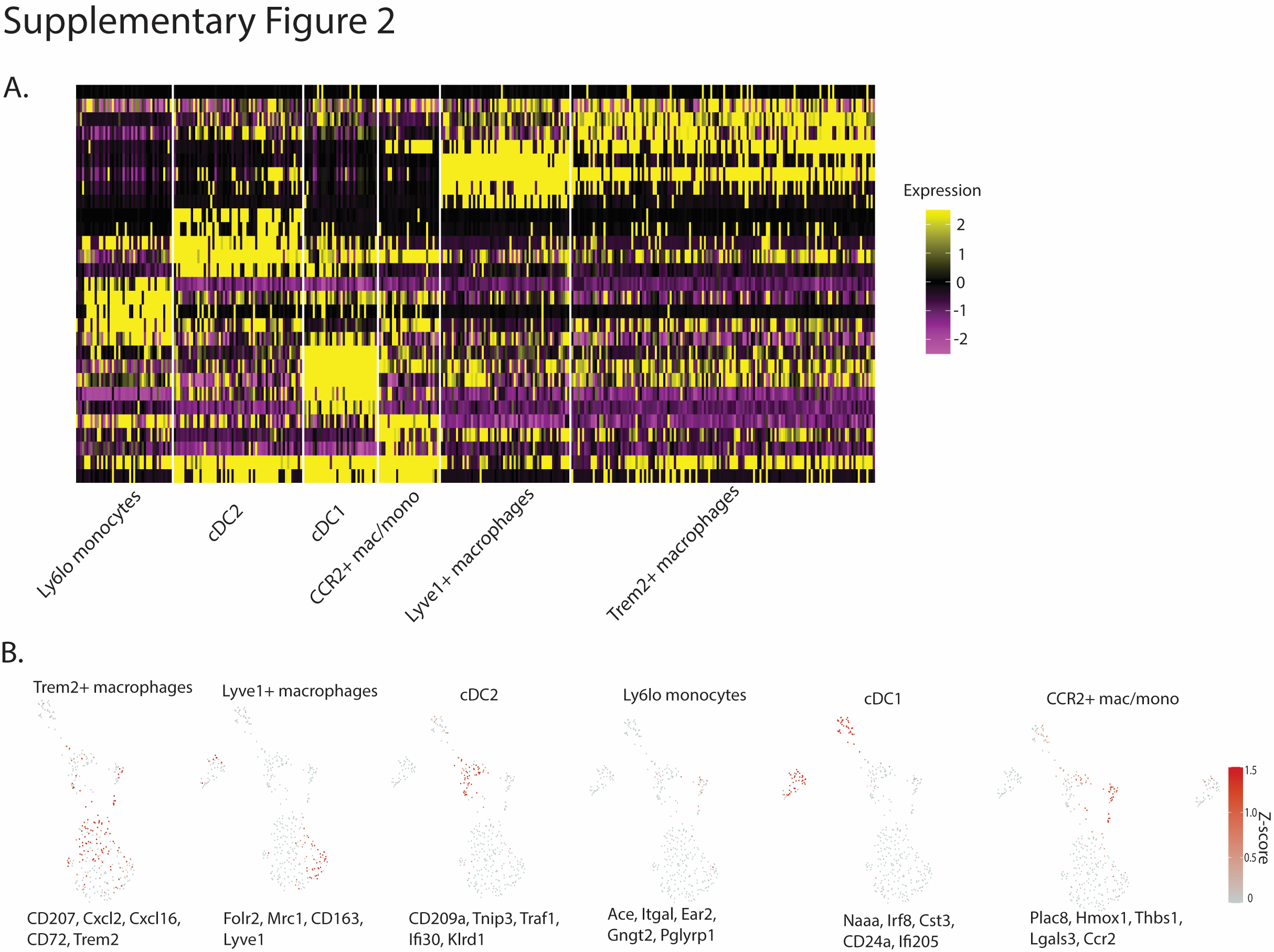

### Supplementary Figure 6.jpg

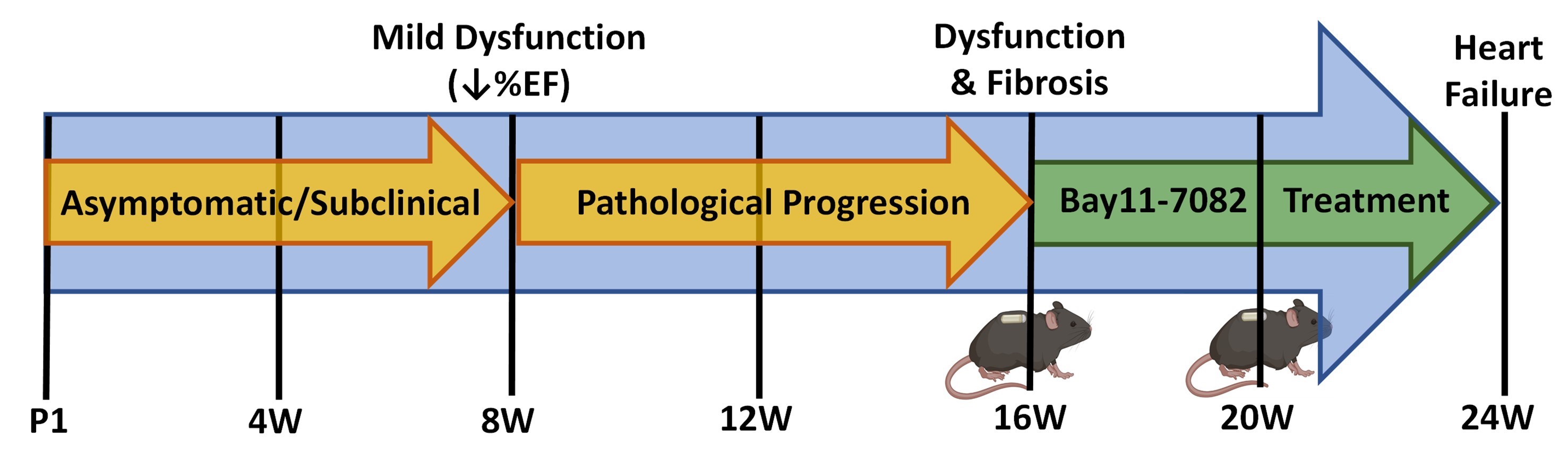

### Supplementary Table 1.jpg

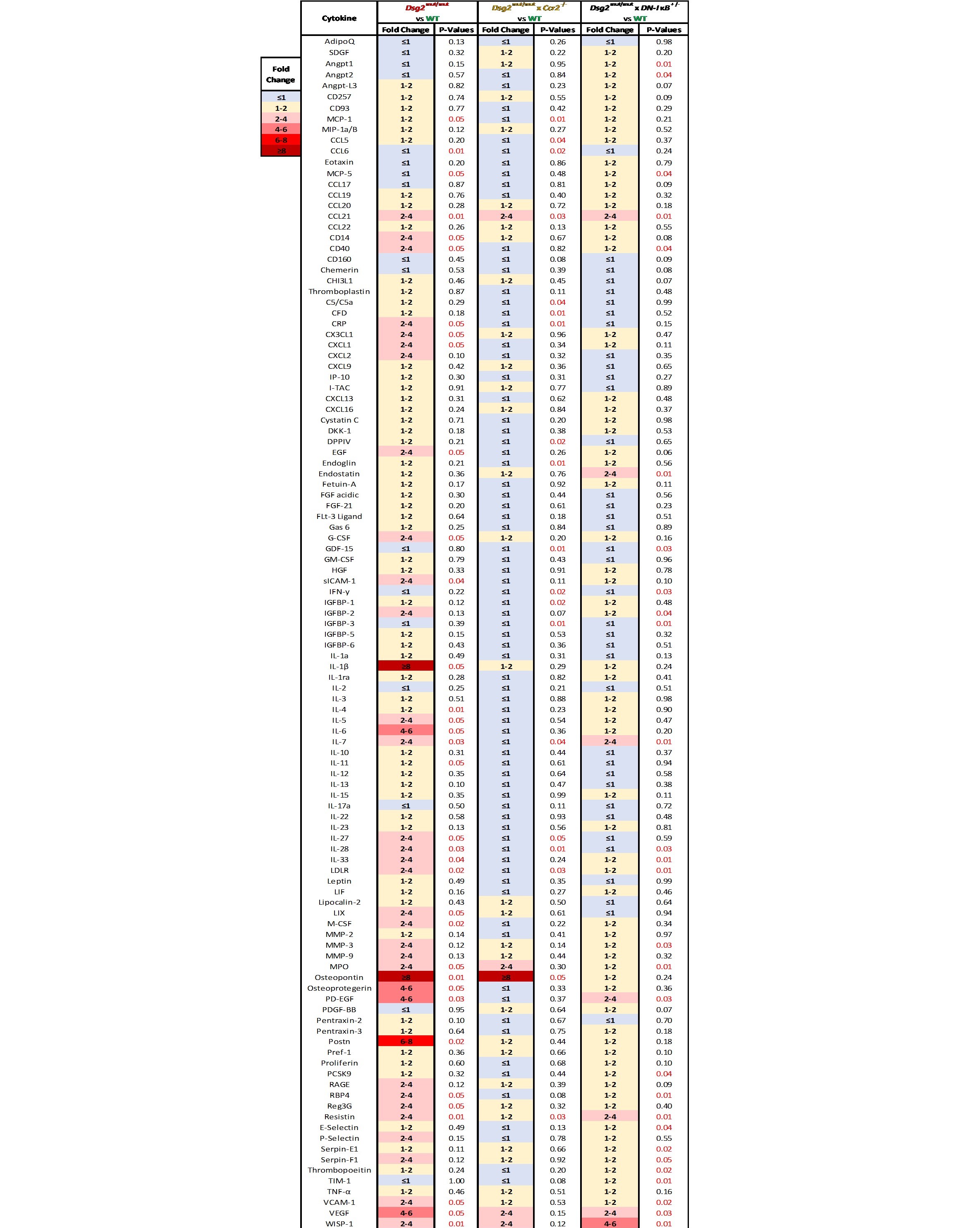
